# Structural insight into binding of novel PET tracer MODAG-005 to lipidic *α*-Synuclein fibrils

**DOI:** 10.1101/2025.04.21.649837

**Authors:** Myeongkyu Kim, Dirk Matthes, Benedikt Frieg, Andrei Leonov, Sergey Ryazanov, Daniel Bleher, Ann-Kathrin Grotegerd, Christian Dienemann, Armin Giese, Gunnar F. Schröder, Stefan Becker, Kristina Herfert, Bert L. de Groot, Loren B. Andreas, Christian Griesinger

## Abstract

Accumulation of α-synuclein (αSyn) aggregates in the human brain is the major hallmark of synucleinopathies such as Parkinson’s Disease, Multiple System Atrophy, and Dementia with Lewy bodies. Positron Emission Tomography (PET) plays a vital role in diagnosing these diseases and monitoring their progression by enabling the non-invasive and sensitive detection of αSyn aggregates in the brain. However, developing PET tracers with specific target binding, as well as identifying the binding site and complex structure present significant challenges. Here, we investigated the interaction between lipidic αSyn aggregates and MODAG-005, a new αSyn PET tracer candidate, using nuclear magnetic resonance spectroscopy, cryogenic electron microscopy and molecular dynamics (MD) simulations. Two binding sites of MODAG-005 were found, one on the surface and one in a tubular cavity of the fibril, of which the occupancies were found to depend on the preparation protocols. The cavity binding site is thermodynamically more stable than the external binding site and is the only site occupied when MODAG-005 is applied with liposome as carriers to the aggregates. This is corroborated by MD simulations in which stable interactions between MODAG-005 and glycine containing backbone motifs of the tubular cavity of the aggregates are observed.

## Introduction

Synucleinopathies are a group of neurodegenerative disorders characterized by pathological accumulation of α-synuclein (αSyn) in brain cells, possibly leading to progressive neuronal dysfunction and ultimately resulting in the loss of nerve cells^1,2^. Prominent examples include Parkinson’s disease (PD)^3^, Dementia with Lewy bodies (DLB)^4,5^, and multiple system atrophy (MSA)^6,7^. αSyn aggregates, which are the major hallmark of these diseases, have become an important target for the development of treatments and diagnostic tools. However, disease-modifying treatment and early diagnosis using PET tracers are not yet available for these diseases^8^, and treatment is limited to alleviating symptoms^9^.

Therefore, accurate and early diagnosis is crucial to assess disease progression and provide molecular-based biomarkers for use in clinical trials. To meet these goals, various diagnostic methods (Dopamine transporter scan, MRI, behavioral measures, seed amplification assays from spinal fluid) are being used^10,11^. Among them Positron Emission Tomography (PET) is a non-invasive imaging technique that traces a radioactively labeled molecule in medical and research settings to visualize and quantify various biological processes in living organisms^12–17^. However, an accurate diagnosis using PET requires a radiotracer with high binding affinity and specificity to disease-specific targets. This includes misfolded and aggregated αSyn, which shows a cell- and region-specific accumulation in synucleinopathies^18^. Several compounds have been tested as potential PET tracers recently, but still show limitations in terms of selectivity and off-target binding^19,20^.

MODAG-005 is a recently developed PET tracer candidate that was developed starting from the molecular structure of emrusolmin (anle138b^21,22^), a small molecule drug targeting αSyn aggregates which is currently tested in a phase II trial (NCT06568237) after successful phase I (Levin et al. 2022; NCT04685265). MODAG-005 detected αSyn aggregates in human MSA brain tissue with very high affinity (K_d_ = 0.2 nM) and specificity, validated through binding assays on recombinant fibrils, microautoradiography and immunohistochemistry^23^. In addition, selectivity to αSyn aggregates was 195-fold over tau and 3600-fold over Aβ aggregates, making it a potential tracer candidate in clinical settings as well as in clinical trials that require longitudinal monitoring of disease progression^23^. However, the binding mechanism for αSyn aggregates is not yet well understood and further information is of high interest with regard to the development of next-generation diagnostic tracers.

We recently reported the segmental folding process of αSyn in the presence of negatively charged liposomes consisting of 1-palmitoyl-2-oleoyl-snglycero-3-phosphate (POPA) and 1-palmitoyl-2-oleoyl-sn-glycero-3-phosphocholine (POPC)^24^ and structures of lipidic αSyn oligomer using NMR spectroscopy, fluorescence microscopy and molecular simulation^25^ as well as lipidic fibrils named L2 fibrils using cryo-electron microscopy (cryo-EM), NMR spectroscopy and molecular simulation^26^. Leveraging the availability of L2 fibrils and their dissociation constants to MODAG-005 - comparable to the high binding affinities observed in autoradiographic studies of brain slices from MSA patients - we here present the binding mode and mechanism of MODAG-005 interaction with lipidic αSyn fibrils (L2) using NMR spectroscopy, cryo-EM, and molecular dynamics (MD) simulations. We first describe the localization of this small organic molecule mainly based on NMR, then show with cryo-EM that the L2 structure is retained irrespective of the presence or absence of MODAG-005 and finally provide insight into the structure of the MODAG-005/L2 complexes based on the experimental results using atomistic MD simulations.

## Results

### MODAG-005 can bind to external and internal sites of lipidic αSyn L2 fibrils

To investigate whether MODAG-005 binds to monomeric αSyn, which would be undesirable for a PET tracer, titration experiments were conducted using solution-state NMR spectroscopy. Overlaid ^1^H-^15^N HSQC spectra of monomeric αSyn only and with 3.5 fold excess (mol/mol) of showed negligible effects on the chemical shifts of aSyn, i.e., negligible binding of MODAG-005 to monomeric αSyn. (Supplementary Fig. 1).

In contrast, binding of MODAG-005 to lipidic αSyn aggregates (L2 fibrils) measured with radioactive MODAG-005, revealed two high affinity binding sites, one with a subnanomolar K_d_ (0.75 ± 0.21 nM) and the second with a K_d_ of 6.06 ± 0.7 nM (Fig 1a, b). The K_d_ of the high-affinity binding site is similar to the one observed when MODAG-005 was tested on brain slices of MSA patients^23^ which suggests that the binding site could be similar to the one in those patient brain fibrils. We determined the binding site of MODAG-005 to L2 fibrils using three different sample preparation conditions. First, exploiting its highly hydrophobic nature, MODAG-005 was added after incorporating it in liposomes to monomeric protein before starting aggregation. Secondly, MODAG-005 was dissolved in dimethyl sulfoxide (DMSO) that was added at a 2% (v/v) ratio to preformed L2 fibrils. The addition of potential PET tracers dissolved in DMSO has been described in several studies for other candidate compounds^27,28^ and was therefore also tested here, although DMSO is a less “physiological” carrier for delivery of lipophilic drugs than liposomes^16,27–30^. Lastly, MODAG-005 was administered with the liposomes to preformed L2 fibrils. These liposomes were small unilamellar vesicles (SUVs) composed of POPA and POPC (1:1) as described before^24,26,31^. Intermolecular interactions between MODAG-005 and SUVs were observed in NMR spectra, indicating the localization of the rather hydrophobic compound within the bilayers of SUVs (Supplementary Fig.2). As a control, hCANH experiments were performed to evaluate the effect of the addition of DMSO and SUVs on L2 fibrils, indicating minimal impact from DMSO or from SUVs (Supplementary Fig.3) on the spectrum of preformed L2 fibrils.

**Fig. 1.**
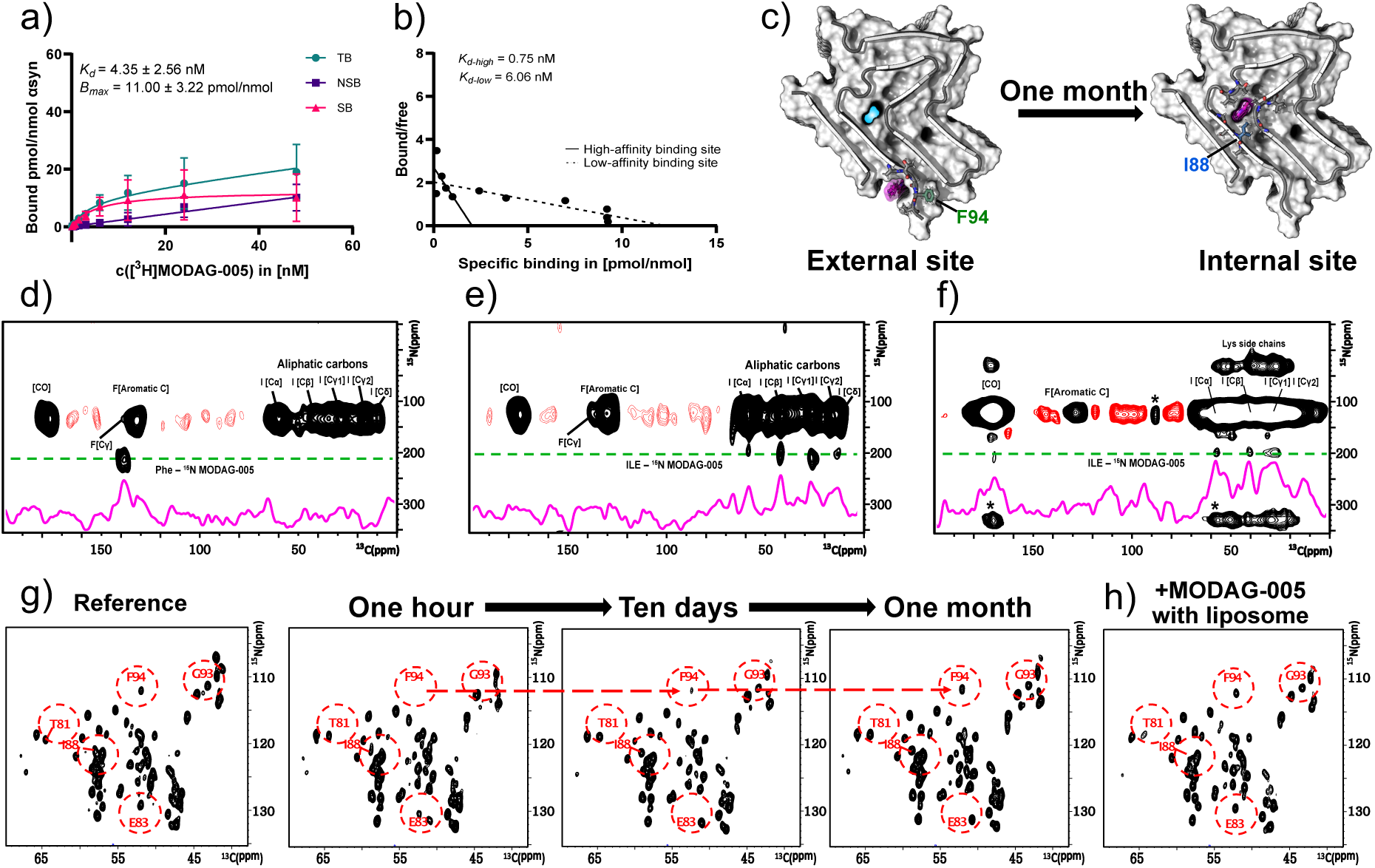
Identification of MODAG-005 binding to *α*Syn fibrils by DNP enhanced MAS NMR. **a)** [3H]MODAG-005 *in vitro* binding experiments. Non-linear regression of TB and NSB revealed a high affinity towards αSyn fibrils (n=4) **b)** Scatchard analysis revealed a high-affinity (dashed line) and a low-affinity (solid line) binding site. **c)** Modelled complex structure of L2 αSyn fibril with MODAG-005 located in external and internal binding site. The spectral region shown contains cross-peaks between pyrazole NH of MODAG-005 (Green line) and protein carbon atoms. 2D NHHC spectra of αSyn fibrils prepared from protein with amino-acid-specific isotope labeling on Ile and Phe-ring. Addition of 15N-labeled MODAG-005 in DMSO **d)** after aggregation and **e)** before aggregation, **f)** 15N-labeled MODAG-005 added in liposomes to L2 fibrils. Proton-proton mixing was 200 µs. **g)** hCANH spectra of L2 with MODAG-005 added in DMSO after fibril formation shows displacement of MODAG-005 from the external to the internal binding site with increasing incubation time: one hour, ten days and one month. **h)** hCANH of complex fibrils in the presence of liposome containing MODAG-005. K_d_, dissociation constant in nM; Bmax, number of accessible binding sites in pmol/nmol. TB, Total Binding; NSB, Non-Specific Binding; SB, Specific Binding. * spinning side bands

The L2 structure of the aSyn fibrils^26,31^ was conserved under all three conditions (vide infra). When MODAG-005 was administered in SUVs during or after αSyn aggregation, MODAG-005 affected the chemical shifts or linewidths of resonances belonging to residues G68, V71, K80, T81, G86, and I88 located around the internal cavity of the L2 fibril structure, similarly to what has been described for anle138b^31^. Only when MODAG-005 was administered with DMSO after aggregation, it had an impact on not only residues forming the internal cavity but also on residues E61, Q62, F94, V95, and K96, which constitute the external site (Fig. 1c, Supplementary Fig. 4). Since chemical shift changes upon binding can be caused by direct contact with the ligand or by allosteric effects induced by the binding, we measured direct contacts between MODAG-005 and the protein. To this end, two-dimensional (2D) NHHC experiments enhanced by dynamic nuclear polarization (DNP) and magic-angle spinning (MAS) at 100 K^32^ were conducted following a published protocol^31^. For that purpose, MODAG-005 was isotopically labeled with ^15^N on its pyrazole nitrogens. Selectively labeled αSyn was expressed in presence of phenylalanine with ^13^C-labeled phenyl ring and ^13^C/^15^N-labeled isoleucine (Fig. 1d-f, Supplementary Fig. 5). It turned out that MODAG-005 binds in the internal cavity near I88 when administered with SUVs irrespective of the time point (during or after aggregation) (Fig 1e,f). When applied in DMSO to the L2 fibrils, MODAG-005 binds to the external binding site near F94. While the DNP spectra do not show binding to the internal binding site for this latter condition, still residues G68 and G86 in the cavity region disappeared from the spectrum, indicating probably too few MODAG-005 molecules in that binding site to be detected by the DNP-NHHC experiment. We cannot exclude allosteric effects through the binding to the external site transmitted to the cavity residues, as well.

In summary, when MODAG-005 was administered with the liposome vehicles, only the internal MODAG-005 binding site is occupied. Application of the compound in DMSO leads to the initial population of an additional, external site.

### The addition of MODAG-005 does not alter the L2 protofilament fold

To investigate the changes in the L2 fibril structure upon the addition of MODAG-005, we employed cryo-EM and solid-state NMR spectroscopy. Three cryo-EM samples were prepared under the same conditions as described for the hCANH experiment. Briefly, the first was L2 fibrils without MODAG-005 as a control, the second was L2 fibrils incubated with MODAG-005 dissolved in DMSO for one hour (final DMSO concentration 2%), and the third was the second sample incubated at 37°C for ten days.

First, solid-state NMR was utilized to analyze the entirety of structures in the sample. we investigated the experimental correlation between carbon atoms by acquiring 2D ^13^C-^13^C correlation spectra using dipole assisted rotational recoupling (DARR)^33^. The 2D ^13^C-^13^C DARR spectra with long mixing (Fig. 2a) show similar protein long-range contacts in the presence and absence of MODAG-005 applied in DMSO, confirming an overall conserved αSyn protofilament fold (Fig. 2b,c). The contact between T92 and V63 in the DARR spectrum of L2 reflects the distance between the two β-strands (β5 & β8, Fig. 2d) in L2, indicating a consistent distance preservation regardless of the presence of MODAG-005.

**Fig. 2.**
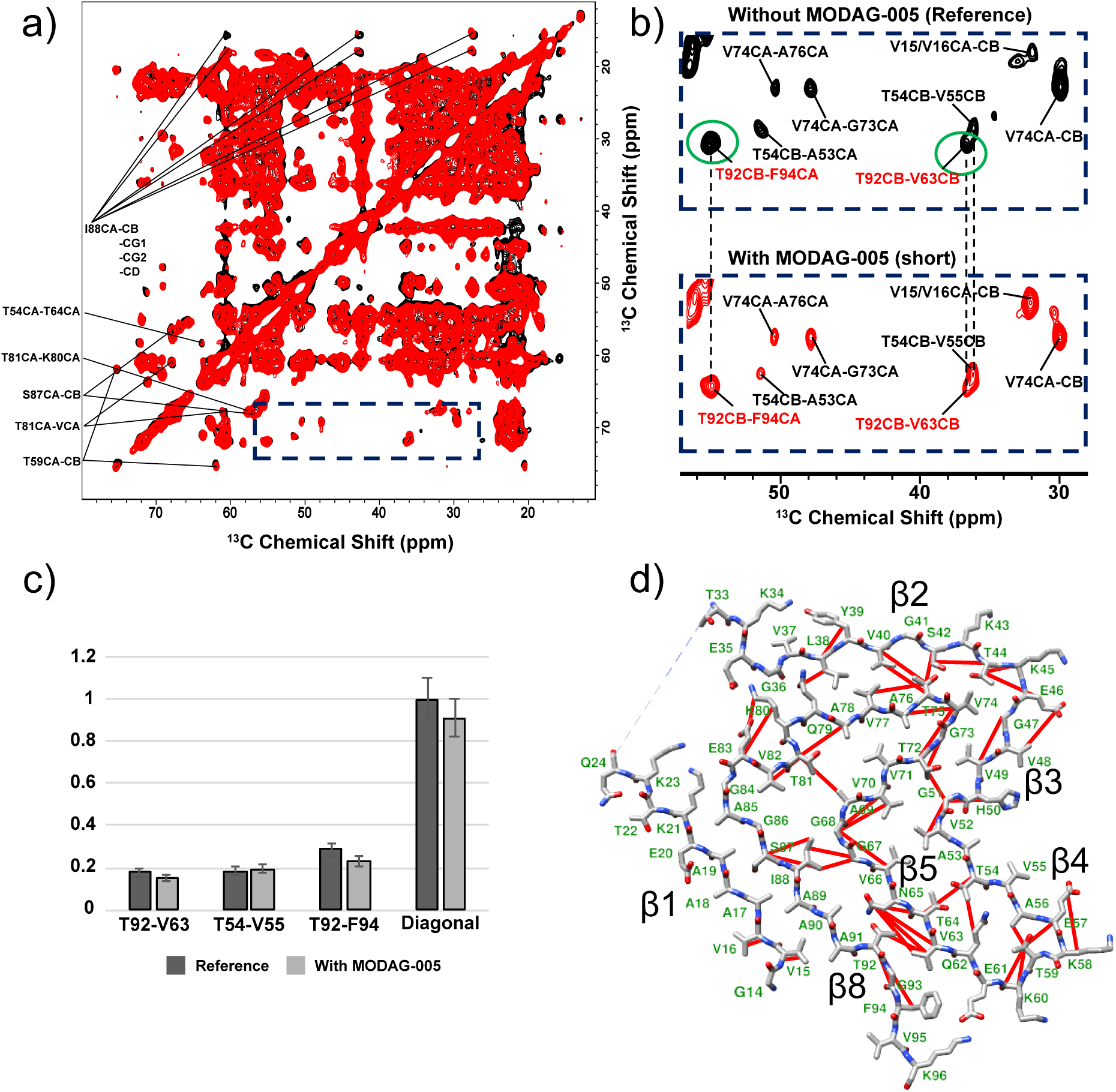
Comparison of the ^13^C-^13^C DARR Spectra of L2. **a)** Overlay of 13C-13C DARR spectra without MODAG-005 (in black, reference) and with MODAG-005 (in red, short) applied in DMSO after fibril formation and short incubation time (one hour at RT). **b)** Zoomed-in view near the F94 position. **c)** Quantitative comparison of the peak heights compared to diagonal peaks. **d)** Carbon-carbon contacts within the L2A structure.

Next, we solved the 3D structures of αSyn in the presence of MODAG-005, using the same batch of fibrils as we used for NMR experiments (Fig. 3). After adding MODAG-005 dissolved in DMSO, fibrils obtained after one hour of incubation (Fig. 3b) and after ten days of incubation at 37°C (Fig. 3c) were compared. Two fibrils with slightly different twist angles in the two collected samples were identified and marked as Ⅰ (-0.739°) and Ⅱ (-0.598°). The twist angle did not affect the L2 protofilament fold, which was the same for all three samples (Supplementary Fig. 7). All protofilaments revealed the L2 fold, which is in agreement with the NMR results, and in addition to the L2A trimeric fibril^26^, we also found an asymmetric dimeric fibril. Altogether, analysis of the cryo-EM and solid-state NMR results confirmed that the structure of L2 remained unchanged, irrespective of whether MODAG-005 was bound externally or internally.

**Fig. 3:**
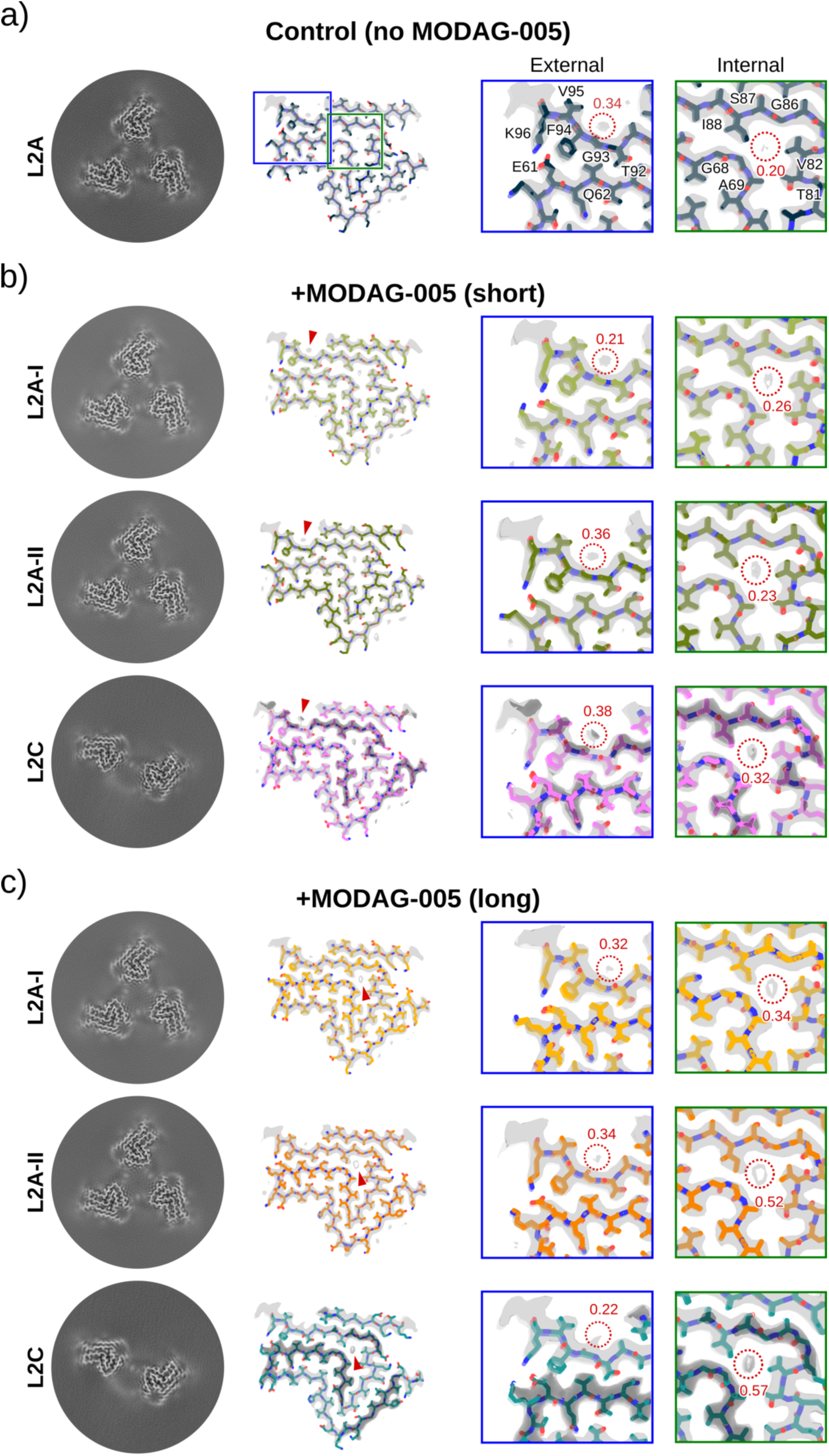
Comparison of complex structures over time. The reference cryo-EM structure of L2A αSyn fibrils (8A4L: Ref. DOI: 10.1038/s41467-022-34552-7) without MODAG-005 is depicted in **a)**, complexes with MODAG-005 added after aggregation (2% DMSO) with short (one hour at room temperature) **b)**, and (Ten days at 37°C) incubation **c)**. From left to right, the panels show a cross-section of the sharpened cryo-EM density map, an overlay of the atomic model (colored stick model) and the sharpened map (transparent surface) of an isolated protofilament with additional densities at the MODAG-005 binding sites indicated by the red triangles, and close-up views of the external and internal MODAG-005 binding sites. In the close-up views, the labels show the average density over a cylinder with a 2.0 Å radius and a height of 4.7 Å (the approximate position is depicted by a red circle). These averaged values have an estimated error of 0.05 and were computed from normalized density maps with a mean of 0.0 and a standard deviation of 1.0.

### MODAG-005 is thermodynamically more stable in the internal cavity than at the external binding site of L2

MODAG-005 binds only to the internal binding site when applied to fibrils via liposomes and this binding site was the only one consistently detected after one hour or one month of incubation (Supplementary Fig. 8a), as inferred from hCANH experiments, which remained unchanged over time. However, when MODAG-005 was added in DMSO to preformed L2 fibrils, we identified an external binding site and subsequently studied the stability of this less “physiological” binding site by varying the incubation time of the sample based on this application protocol. When we observed the hCANH spectra of samples incubated for one hour, ten days, or one month, the NMR signals of residues F94, G93, and V95 gradually reverted to the characteristic L2 state without MODAG-005, indicating that MODAG-005 left the external binding site after ten days (Fig. 1g). Conversely, residue I88 in the internal cavity was also affected by MODAG-005, but its chemical shift remained consistent with the spectrum observed when MODAG-005 was initially administered in DMSO. In contrast, residue E83 showed a slightly different spectrum than initially, suggesting that E83 is located in the hinge region of the inner cavity, where the flexibility may change slightly as MODAG-005 molecules populate the cavity region of the fibril (Supplementary Fig. 8b). These findings demonstrate that the MODAG-005 binding is more stable in the internal site compared to the external site, ultimately leading to the population shift of MODAG-005 from the external to the internal binding site (Fig. 1g, Supplementary Fig. 9).

Next, we analyzed the cryo-EM maps of the MODAG-005/L2 structures obtained immediately after addition of MODAG-005 in DMSO and ten days later. All cryo-EM maps show additional densities that are not connected to the protein density, in particular in the tubular cavity formed by the residues G67-V70 and T81-I88.

We calculated the average density for this internal site and compared it to the L2A fibrils in the absence of MODAG-005. The average density is higher in the presence of MODAG-005 compared to the L2A fibrils, and also increases with incubation time with MODAG-005 (Fig. 3b, c). Combined with the insights from NMR experiments, this data suggests that the additional density is related to the position of MODAG-005 bound to the αSyn fibril, although the resolution of the cryo-EM density does not allow a more detailed interpretation.

NMR experiments revealed MODAG-005 interacting with E61, Q62, F94, V95, and K96 at the L2 fibril surface. We found only weak non-fibrillar densities close to F94 in all fibrils. As a similar non-fibrillar density is also found in the reference cryo-EM L2A structure, the additional density in our fibril in the present study cannot be unambiguously assigned to MODAG-005 bound to the fibril surface. A plausible explanation is that the external binding site is easily accessible to MODAG-005, such that it binds to that particular site with high off-rate, which would make structure determination quite challenging. In addition, MODAG-005 may bind irregularly, non-equally spaced along the helical axis to the αSyn fibril, which makes the detection by cryo-EM even more challenging.

It is also interesting to note that we found two different populations of L2A fibrils, for short and long incubation time with MODAG-005, which only differ in slight variations of the twisting angle between the stacked peptides along the helical axis. Whether this is an effect of MODAG-005 cannot be answered by our data conclusively.

### Backbone interactions to or near glycine residues dominate the binding mode of MODAG-005 within the internal and external cavities of L2 fibrils

While NMR provided experimental distances between the pyrazole nitrogens and amide protons of the L2 and cryo-EM provided the L2 structure in complex with MODAG-005 that was identical to the PDB entry of the free L2 form (PDB ID 8A4L)^26^, we continued to build an L2 model with bound conformations of MODAG-005 modelled within the internal and external binding sites as initial structure for fully atomistic MD simulations αSyn (see “Methods” and Supplementary Table 6). The L2 model protofilament structure is composed of 30 αSyn molecules and was simulated with position restraints on the backbone atoms of the residues resolved by cryo-EM (Supplementary Fig. 10). We extended the model structure with a flexible N- and C-terminus from other published αSyn polymorphs such that the putative binding pocket around G93F94 is better defined (Fig. 4a, compare a with b and c in Supplementary Fig. 10). Due to the dynamic nature of the N- and C-terminus, these parts are not visible in the L2 cryo-EM structure (Supplementary Fig. 10c,d). We then simulated the fibril model in the presence of POPC and POPA lipids and recorded the distribution of minimum distances between the pyrazole nitrogen atoms of MODAG-005 and amino acid residues in both the internal and external binding site of L2 (Fig. 4b). Polar contacts between MODAG-005 atoms from the amino group of the N-methylaniline, the nitrogen atoms of the pyrazole, as well as the bromine atom of the bromopyridine and protein backbone atoms were observed within the internal cavity to residues I88, G67, and G68 (Fig. 4c-f). In addition, the long axis of MODAG-005 was aligned with the protofilament axis and the molecule exhibited discrete translational motion along this axis (Fig. 4c, Supplementary Fig. 11). For the external binding site polar contacts were found between MODAG-005 and protein backbone atoms of residues A91-F94 (Fig. 4c-f).

**Fig 4.**
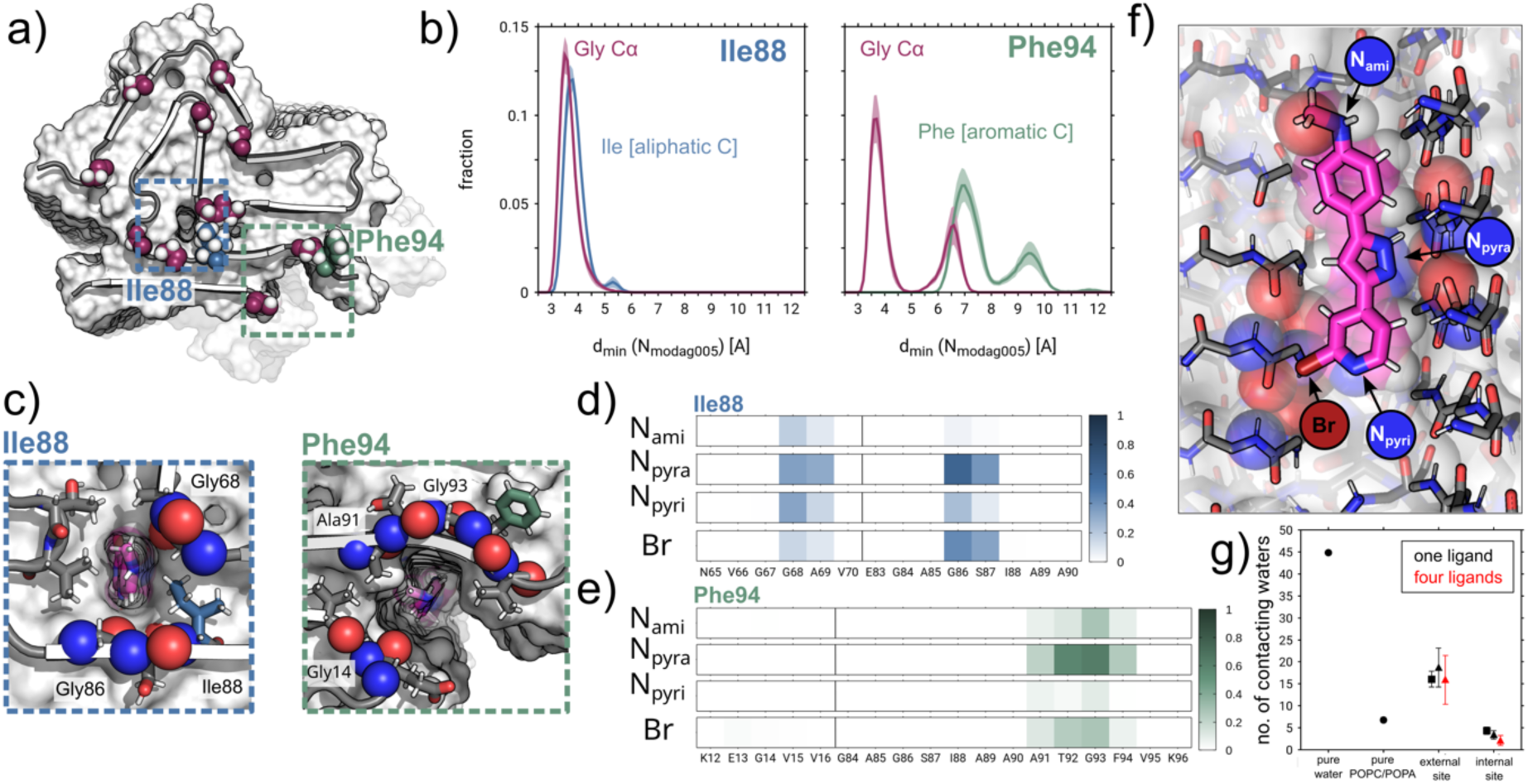
MD simulations recapitulate interatomic distances for the two putative MODAG-005 *α*Syn fibril binding sites. **a)** Starting structure for MD simulations and location of the two putative MODAG-005 binding sites: In the internal cavity (light blue box) of αSyn protofilament L2 between residues 67GGAV70 and 80KTVEGAGSI88 or in an external cavity (light green box) between residues 14GVV16 and 91ATGFV95 (shown without MODAG-005 for clarity). The rigid part of the αSyn protofilament L2 cryo-EM structure (residues 14-24, 33-96) is shown in cartoon representation and with color-coded residues: Gly (magenta), Ile88 (blue), and Phe94 (green). The N- and C-terminally extended model of L2 used for the MD simulations (residues 10-24, 33-100) is shown in surface representation in the background. **b)** Distributions of minimal distances between pyrazole nitrogen atoms of MODAG-005 and the aliphatic carbon atoms of Ile88 (light blue), and Cα of G67, G68, G84, G86 (magenta) for the internal binding site. Distributions of minimal distances between pyrazole nitrogen atoms of MODAG-005 and the aromatic carbon atoms of Phe94 (light green), and Cα of G14 and G93 (magenta) for the external binding site. **c)** Tubular cavities of internal and external binding site with bound MODAG-005 viewed up close and down the long axis of the αSyn protofilament. Prominent MODAG-005 contact sites in the backbone scaffold are represented by spheres (blue - amide nitrogen; red - carbonyl oxygen). **d)**, **e)** Backbone atom contacts of individual residues with polar moieties of MODAG-005 as defined in **f)**. Scale bars (right) indicate contact probabilities. g) Average number of water oxygen atoms in direct contact with the MODAG-005 ligand in different simulation scenarios and binding sites. Black and red symbols denote simulations with one or four MODAG-005 ligands simultaneously present. Circles report simulations without fibril models, squares report data for simulations of full-length L2 protofilament model and triangles for truncated fibril model systems.

The patterns of polar protein and ligand interactions were reproduced by MD simulations of model systems of the putative small molecule binding sites where the MODAG-005 ligand was initially placed outside of the fibril (Supplementary Fig. 12 and 13). The simulations suggest that MODAG-005 can bind in multiple degenerate poses with different ring orientations to the backbone of the respective residues surrounding the binding sites (internal: G68A69; G86-I88 and external: E13-V15; A91-V95). MM/PBSA based energetic analysis of the MD simulation data confer a relatively higher stability of MODAG-005 binding to the internal compared to the external binding site (Supplementary Fig. 14 and Supplementary Table 7). We determined the average number of water oxygen atoms in direct contact with the MODAG-005 ligand in the two different binding sites, as well as water and lipids as local environment. The external MODAG-005 binding pose displayed a higher degree of water solvation compared to the internal binding site, or MODAG-005 simulated in a pure mixture of POPC/POPA lipids. This observation provides insight into the degree of solvent exposure of MODAG-005 at the fibril exterior as a driving force for its potential reversible unbinding from the external binding site. Additionally, MD simulations were started with multiple copies of the ligand. These simulations indicate the ability of MODAG-005 to transiently form tight stacks/bunches of vertically bound molecules inside the internal cavity of L2 (Supplementary Fig. 13). This is attributed to close interactions between the ligands within the internal cavity, specifically between the amide group on one side of a MODAG-005 molecule and the pyrimidine nitrogen and bromine on the other side of an adjacent MODAG-005 molecule.

## Discussion

Clinical assessment of amyloid burden through PET imaging has played a crucial role in the outcome evaluation of clinical trials to investigate the effectiveness of anti-amyloid treatments^34,35^. In the past, several PET radiotracers have been developed for the purpose of imaging neuropathological features associated with AD and tauopathies^36–38^. However, to date, developing tracers that bind effectively to αSyn aggregates in the human brain has proven difficult due to insufficient binding affinity, low selectivity and off-target binding^27,39–42^ For example, the probes SH-299, SL-631, LDS-798, Nile Red, and Nile Blue, which are based on Thioflavin-T or Thioflavin-S, both are non-selective amyloid dyes, were studied but did not advance to PET development due to low binding or poor selectivity^43,44^. However, they were used as lead structures for developing alternative αSyn selective ligands.

Several research groups are trying to understand the binding mechanism of candidate molecules to various protofilament structures. Representative examples^45^ include compounds such as MODAG-005^23^ related to anle138b^21,22,46^, ACI-12589^19^, F0502B^28^, 2FBox^40^, 15a^47^, S3-1^48^, C05-05^20^, SPAL-T-06^49^ and TZ6184^50^. Among them, the candidates with high affinity (K_d_ less than 1 nM) for αSyn aggregates are MODAG-005 and TZ6184. TZ6184 has an exceptionally good K_d_ value of 0.39 nM, but its selectivity profile has not been reported, and only limited information is available. For MODAG-005 detailed in-vitro and in-vivo evaluation results are available^23^.

So far, tracer targets have primarily been identified at external binding sites, similar to those observed in fibrils of HET-s^51^, Aβ^52^, and tau^30,53^. However, both theoretical studies^54^ and experimental data^31,55^ suggest that small molecules can also be situated in the αSyn fibrillar cavity. Notably, for anle138b selective binding was detected basically in the same internal cavity as identified for MODAG-005, along with dynamic movement of anle138b along the fibril axis due to the translational symmetry of the fibril (Supplementary Fig. 4).

In the present study, we found that the dominant binding site of MODAG-005 is the internal binding site when MODAG-005 was applied in the presence of lipids irrespective of exposure time. This is also the situation in which a PET tracer is applied *in vivo*. It is tempting to compare the expected off-rate with the wash out of MODAG-005^23^. Assuming a reasonable on-rate of 10^6^/(Ms) for MODAG-005, an off rate of 22 min would result for the measured K_d_ of 0.75 nM which is in the ball park of observation for the brain PET. By contrast, only when MODAG-005 was applied with DMSO to preformed lipidic fibrils of type L2 without an excess of free lipid molecules, the external binding site was populated. Yet, the thermodynamic sink is the internal binding site, since with time MODAG-005 migrated away from the external binding site.

Our results characterize the binding site of MODAG-005 in the L2 fibrils and provide a model for the mechanism by which the PET tracer candidate binds to specific sites in the fibril when lipids are included in the preparation. Although L2 is an in vitro fibril, it mimics a physiological brain environment by incorporating lipids. The L2 fibrils are composed of a hydrophobic core with a fold similar to previously reported patient-derived or seeded fibrils (PDB: 8A9L, 6XYQ, 8CYT, 7OZG, 7NCH and 7XO2), featuring an internal cavity. Although L2 has limitations in representing polymorphs without an internal cavity (PDB: 7OZG), it represents polymorphs with a hydrophobic core in the internal cavity quite well, as they are structurally similar. This hydrophobic core portion is a binding target for potent fibril binding compounds (e.g., anle138b and Pittsburgh Compound B^36^), further rationalizing the favorable MODAG-005 binding to the tubular cavity of the L2 fibril (Supplementary Fig. 14 and Supplementary Table 7).

Based on this model, we expect to predict and utilize specific binding sites for effective PET tracers and therapeutic drugs for different αSyn folds from other disease-associated MSA and DLB αSyn fibril structures with internal cavities (PDB ID 8A4L and 6XYQ) (Supplementary Fig. 14 and Supplementary Table 7), as well as amyloidogenic proteins such as Aβ, tau, and prion protein.

## Methods

### Protein expression and purification

αSyn was expressed recombinantly in E. coli strain BL21(DE3) and purified as described previously^56^. Briefly, the protein was expressed in minimal medium at 37 °C. Cells were harvested 6 h after induction, lysed by freeze-thaw cycles followed by sonication, boiled for 15 min and centrifuged at 48,000 × g for 45 min. From the supernatant DNA was precipitated with streptomycin (10 mg/ml) while stirring the ice-cold solution. After centrifugation αSyn was precipitated from the supernatant by adding ammonium sulfate to 0.36 g/ml. After another centrifugation step the pellet was resuspended in 25 mM Tris/HCl, pH 7.7 and the protein was further purified by anion exchange chromatography on a 30 ml POROS HQ column (PerSeptive Biosystems). To prepare monomeric αSyn without any aggregates, the protein was dialyzed against PBS buffer, pH 7.4, centrifuged at 106,000 × g for 1 h at 4 °C and filtrated through 0.22 µm ULTRAFREE-MC centrifugal filter units (Merck Millipore). The final protein concentration was adjusted to 0.33 mM.

### Preparation of isotope labeled protein

Uniformly ^15^N- and ^13^C,^15^N-labeled samples were expressed in minimal medium supplemented with ^15^NH_4_Cl and ^13^C_6_-D-glucose (Cambridge Isotope Laboratories and Sigma Aldrich). For production of amino acid specific forward-labeled protein (^13^C-ring Phe, ^13^C,^15^N-Phe, ^13^C,^15^N-Ile), the labeled amino acids were added to the minimal medium 1 hour before induction of protein expression. The protein was finally dialyzed against buffer (50 mM HEPES, 100 mM NaCl, pH 7.4) to obtain a 0.3 mM solution and the resulting solution was stored at -80 °C until use.

### Preparation of isotope labeled MODAG-005

The General Information for the synthesis of compounds (7a = MODAG-005) was described previously^39^.

### Preparation of 4-[5-(2-bromopyridin-4-yl)-1*H*-[1,2-^15^N_2_]-pyrazol-3-yl]-*N*-methylaniline ^15^N_2_-MODAG-005

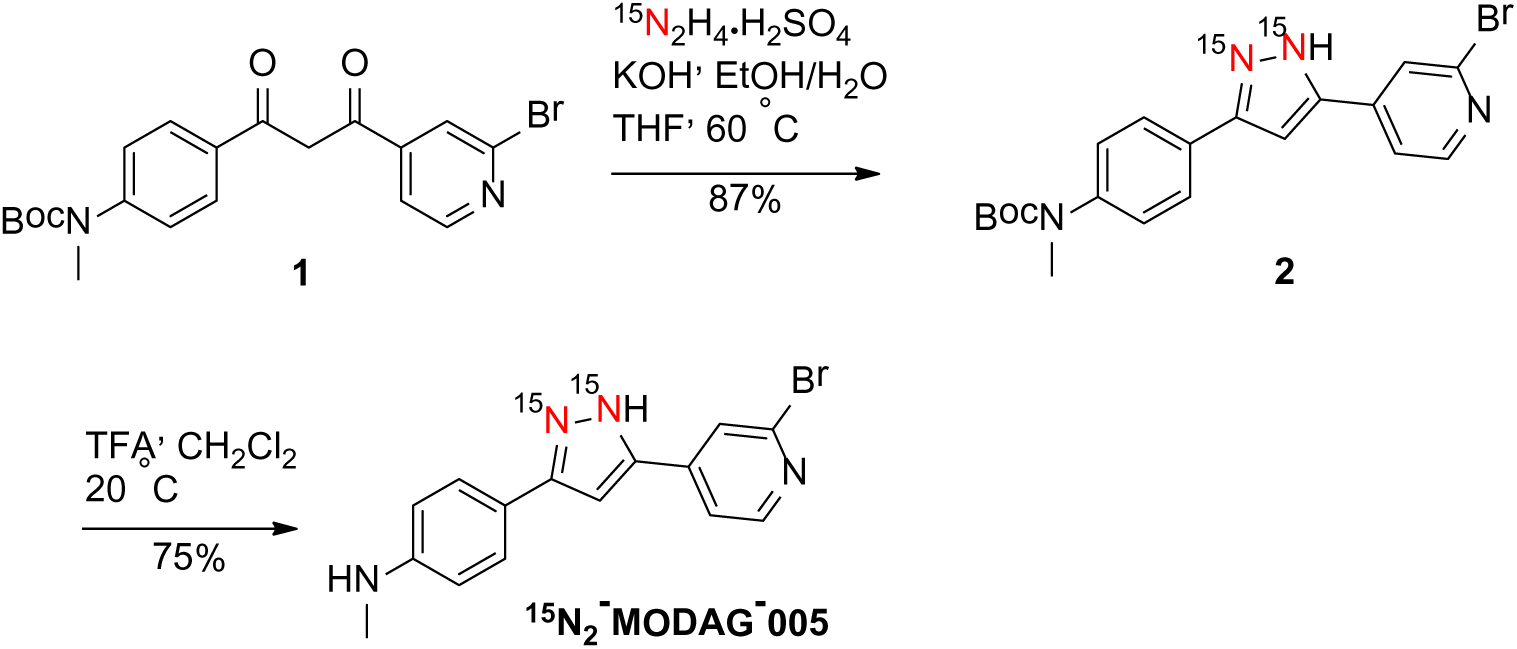

A mixture of [^15^N_2_] hydrazine sulfate (1.32 g, 10.0 mmol), 99.5% EtOH (10 mL), water (1.0 mL) and 40% KOH aqueous solution (2.95 g, 21 mmol) was stirred in a sealed pressure flask at 80 °C for 1 h. The mixture was cooled, the solid was filtered off (syringe filter) and rinsed with EtOH (1 mL). The combined filtrate was added to a solution of *tert*-butyl 4-[3-(2-bromopyridin-4-yl)-3-oxopropanoyl]phenyl methylcarbamate **1**^39^ (2.17 g, 5.0 mmol) in THF (25 mL). The reaction mixture was stirred at 60 °C for 24 h, cooled to room temperature, concentrated *in vacuo* and evaporated with methanol (10 mL). The residue was purified by flash chromatography on silica gel (CHCl_3_/MeOH = 95/5, R_f_ = 0.31) to give the intermediate compound **2** (1.88 g, 4.35 mmol, 87%), the protected product as a beige foam. It was dissolved in in CH_2_Cl_2_ (20 mL), trifluoroacetic acid (4 mL, 5.92 g, 52 mmol) was added. The mixture was stirred at room temperature for 15 h and concentrated *in vacuo*. 1M phosphate buffer pH 7 (20 mL) was added, stirred for 1 h, the resulting precipitate was filtered off, washed with water and air dried for 15 h to give the crude product (1.4 g). Crystallization from *n*BuOH (25 mL) afforded the title product, 4-[5-(2-bromopyridin-4-yl)-1*H*-[1,2-^15^N_2_]-pyrazol-3-yl]-*N*-methylaniline 1.08 g (3.26 mmol, 75%) as a beige-colored solid. ^1^H NMR (400 MHz, DMSO-d_6_ + 0.5% TFA) *d* = 8.44 (d, *J* = 5.1 Hz, 1H), 8.08 (d, *J* = 0.8 Hz, 1H), 7.98 (d, *J* = 8.7 Hz, 2H), 7.89 (dd, *J* = 5.1, 1.4 Hz, 1H), 7.66 (d, *J* = 8.7 Hz, 2H), 7.57 (dd, *J* = 3.7, 2.4 Hz, 1H), 2.92 (s, 3H). ^13^C NMR (100.6 MHz, DMSO-d_6_ + 0.5% TFA) *d* = 150.9, 145.6 (2C), 143.2, 142.9, 142.3, 126.4 (2C), 124.3, 123.1, 119.1, 117.9 (2C), 100.9, 32.9.

LC MS (RP18-100Å, gradient 5% CH_3_CN /95% H_2_O to 100% CH_3_CN in 30 min), RT 16.2 min and mass 331.1 (100%), 333.2 (98%), [M+H]^+^. TLC (SiO_2_, *n*-hexane/EtOAc = 1/1) R_f_ 0.4, m.p. 225-226 °C.

### Preparation of αSyn fibrils

αSyn fibrils were prepared as previously reported^24^. In brief, vesicles were prepared by mixing 1-palmitoyl-2-oleoyl-*sn*-glycero-3-phosphocholine (POPC), 1-palmitoyl-2-oleoyl-*sn*-glycero-3-phosphate (POPA, sodium salt) and MODAG-005 dissolved in chloroform respectively and evaporating the solvent under a N_2_-stream and lyophilizing overnight. SUVs were obtained by repeated sonication of a solution of 1.5 mM POPC, 1.5 mM POPA and 150 μM MODAG-005 (Supplementary Table 1). Vesicles were incubated with 70 μM αSyn in buffer (50 mM HEPES, 100 mM NaCl, pH 7.4) at a lipid to protein ratio of 10:1 and a compound to protein ratio of 1:2 and subjected to repeated cycles of 30 s sonication (20 kHz) at 37 °C followed by an incubation period of 30 min. After 96 h fibrils were harvested by centrifuging at 55,000 rpm (TLA-100.3 rotor in an Optima™ MAX-TL) for 1 hour at 4 °C. The supernatant was removed, and centrifuged again 65,000 rpm for 10 min at 18 °C. Excess moisture was removed, and the pellet was immediately packed into 1.3 mm and 3.2 mm rotors for ssNMR. All samples containing MODAG-005 were prepared with the same protein/small molecule ratio.

### Solid state NMR spectroscopy

For ssNMR studies, three types of samples were prepared (see Supplementary Table 2). Fibrils were grown from ^2^H-^13^C-^15^N-αSyn in the presence of vesicles of POPA and POPC (1:1) with or without MODAG-005 at a L/P ratio of 10:1. Fibrils were packed into ssNMR rotors by cutting off the bottom of the tube and centrifuging the pellet directly into the rotor of choice through a custom-made filling device made from a truncated pipette tip. Last, the sample was centrifuged into the rotor in an ultracentrifuge packing device for 10 min at 16,873 × g in a F-45-18-11 Rotor in a 5418 R tabletop centrifuge (Eppendorf, Hamburg, GER).

3D (H)CANH experiments^57^ on fibrils in the presence and absence of MODAG-005 were recorded on an 800 MHz Bruker Avance III HD spectrometer at a magnetic field of 18.8 T equipped with a 1.3 mm magic-angle spinning (MAS) HCN probe and MAS at 55 kHz. The temperature of the cooling gas was set to 250 K, resulting in an estimated sample temperature of 20 °C.

3D (H)CANH spectra on fibrils in the absence and presence of MODAG-005 were recorded on a 800 MHz Bruker Avance III spectrometer at a magnetic field of 18.8 T equipped with a 1.3 mm HCN probe and MAS at 55 kHz. The temperature of the cooling gas was set to 250 K, resulting in an estimated sample temperature of 20 °C.

2D ^13^C^13^C-DARR spectra with and without MODAG-005 were acquired on an 850 MHz Avance III spectrometer with a 3.2-mm MAS HCN probe with mixing times of 20-200 ms at a magnetic field of 19.97 T and MAS at 17 kHz. The temperature of the cooling gas was set to 265 K, resulting in an estimated sample temperature of 20 °C.

3D (H)CANH spectra were acquired in three blocks of 20.5 h in the presence and in the absence of MODAG-005. All 2D ^13^C^13^C DARR spectra were acquired in 4 blocks of 18 h in the presence and 6 equivalent blocks in the presence of MODAG-005. All related blocks were corrected for linear drift of the static magnetic field using an in-house program executed from the command line in Bruker Topspin^58^. The drift corrected blocks were then averaged and processed as one spectrum. Spectra were analyzed using CcpNmr Analysis and NMRFAM-Sparky^59^. Assignments on αSyn fibrils in the absence of MODAG-005 had previously been reported^24^.

### DNP enhanced ssNMR

For DNP enhanced ssNMR, fibrils were grown as for ssNMR measurements, but with isotopic labeling (see the ‘Preparation of isotope labeled protein and isotope labeled MODAG-005’ section) as indicated in the text.

TEMTriPol-1^60^ was synthesized based on published protocols^61^. A solution of ^13^C-depleted *d_8_*-Glycerol (70%-vol in water), containing 8 mM TEMTriPol was added to the resulting aggregate-lipid mixture and mixed thoroughly. Samples were packed into 3.2 mm zirconia ssNMR rotors via a custom-made filling device made from a truncated pipette tip and the rotors were directly frozen in liquid nitrogen before starting the experiment.

All DNP spectra were recorded on a 600 MHz Bruker Avance III HD spectrometer at a magnetic field of 14.1 T equipped with a 3.2 mm low temperature (LT) HCN MAS probe. Detail of the sample conditions are indicated in Supplementary Table 3. For DNP radical-proton transfers 395 GHz microwave irradiation from a gyrotron oscillator was delivered to the sample through a corrugated waveguide. Samples were cooled with a second-generation BRUKER liquid nitrogen cold cabinet, operating at approx. 100 K.

The DNP ssNMR experiments were performed with a recycle delay of 2-4 s (1.2 x T_1_) for each sample and radio frequency power levels of 100 kHz, 72 kHz and 42 kHz for ^1^H, ^13^C and ^15^N respectively.

2D ^13^C-^13^C spectrum 1024 increments were acquired using 1.3 ms RFDR mixing using an xy-8 phase cycling and 83 kHz pi-pulses. ^1^H-^13^C cross-polarization (CP) was applied for 1.5 ms with a 90-100 % ramp on the proton channel.

For NHHC spectra 10 increments (adding MODAG-005 in the presence of liposome: 20 increments) were acquired with 200 μs of proton-proton mixing. The ^1^H-^13^C CP was set to 300 μs to limit transfers to the direct proton environment and ^1^H-^15^N CP was applied for 800 μs with an 80-100 % ramp on the proton channel.

### Solution state NMR spectroscopy

2D ^1^H-^15^N HSQC spectrums were acquired with 32 scans using 256 increments in the indirect dimension and the temperature was set at 278 K. For determination of the conformation of MODAG-005 in the presence of lipids, we prepared a 3 mM solution of vesicles consisting of POPC and POPA (1:1) containing 150 μM of MODAG-005 in buffer (50 mM HEPES, 100 mM NaCl, pH 7.4) with 10% D_2_O. NOESY spectra were acquired with 64 scans using 512 increments in the indirect dimension and a relaxation delay of 1 s. The mixing time was set to 250 ms. Proton 90° flip pulses were 7.4 μs. Experiments were recorded on a Bruker 800-MHz spectrometer. Temperature during measurements was kept at 298 K. 2D datasets were processed in NMRPipe^62^ and analyzed in NMRFAM-Sparky.

### Cryo-EM sample preparation

All cryo-EM samples were identical to NMR samples. Concentration of cryo-EM samples were optimized after initial screening by TEM. To concentrate the fibrils, samples were centrifuged for amples were centrifuged for 5 min at 16,873 × g in a F-45-18-11 Rotor in a 5418 R tabletop centrifuge (Eppendorf, Hamburg, GER). If fibrils did not pellet right away, the procedure was repeated until a visible pellet was obtained. The supernatant was removed and 50 μL of fresh buffer (5 mM HEPES, pH 7.4) were added and thoroughly mixed with the pellet to obtain a highly concentrated fibril solution.

### Cryo-EM grid preparation and imaging

For cryo-EM grid preparation, 1.5 µL of fibril solution were applied to freshly glow-discharged R2/1 holey carbon film grids (Quantifoil). After the grids were blotted for 12 s at a blot force of 10, the grids were flash frozen in liquid ethane using a Mark IV Vitrobot (Thermo Fisher) operated at 4 °C and 95% rH.

Cryo-EM datasets were collected on a Titan Krios transmission-electron microscope (Thermo Fisher) operated at 300 keV accelerating voltage and a nominal magnification of 81,000× using a K3 direct electron detector (Gatan) in non-super-resolution counting mode, corresponding to a calibrated pixel size of 1.05 Å. Data was collected in EFTEM mode using a Quantum LS energy filter at a slit width of 20 eV. A total of 17,071 movies were collected with SerialEM^63^. Movies of were recorded over 40 frames accumulating a total dose of ∼40 e^-^/A^2^. The range of defocus values collected spans from −0.6 to −2.3 μm. Collected movies were motion corrected and dose weighted on-the-fly using Warp^64^.

### Helical reconstruction of αSyn fibrils

αSyn fibrils in complex with MODAG-005 were reconstructed using RELION-3.1^65^, following the helical reconstruction scheme^66^. Firstly, the estimation of contrast transfer function parameters for each motion-corrected micrograph was performed using CTFFIND4^67^. Next, filament picking was done using crYOLO^68^.

The reconstruction procedure was adapted from the original lipidic L2 αSyn fibril^26^. Hence, for 2D classification, we extracted particle segments using a box size of 600 pix (1.05 Å/pix) downscaled to 200 pix (3.15 Å/pix) and an inter-box distance of 13 pix. We separated L2A from L2C fibril by visual inspection of the class images and manual assignment.

For 3D classification, the classified segments after 2D classification were (re-)extracted using a box size of 250 pix (1.05 Å/pix) and without downscaling. Starting from featureless cylinder filtered to 60 Å, several rounds of refinements were performed while progressively increasing the reference model’s resolution. The helical rise was initially set to 4.75 Å and the twist was estimated from the micrographs. Once the β-strands were separated along the helical axis, we optimized the helical parameters (final parameters are reported in Supplementary Table 4). During local refinement, L2A-I and L2B-II finally separated into two individual classes, which were treated individually hereafter.

We then performed a gold-standard 3D auto-refinement, followed by standard RELION post-processing with a soft-edged solvent mask that includes the central 10 % of the box height yielded post-processed maps (B-factors are reported in Supplementary Table 4). The resolution was estimated from the value of the FSC curve for two independently refined half-maps at 0.143 (Supplementary Fig. 6). The optimized helical geometry was then applied to the post-processed maps yielding the final maps used for model building.

### Atomic model building and refinement

For the αSyn fibril, one protein chain was extracted from PDB-ID 8A4L^26^. Subsequent refinement in real space was conducted using PHENIX^69^ and Coot^70^ in an iterative manner. The resulting models were validated with MolProbity^71^ and details about the atomic models are described in Supplementary Table 5.

### Fibril binding assays

MODAG-005 was tritiated by unspecific tritiation via hydrogen exchange with [^3^H]H2 gas (RC Tritec AG, Teufen, Switzerland) resulting in an activity concentration of 1 mCi/mL, >99 % radiochemical purity, and a molar activity of 90.6 Ci/mmol. For saturation binding assays, an optimal fibril concentration of 50 nM for the saturation assay was determined according to Auld et al.^72^ using a concentration determination assay. A 1:2 dilution series of the [^3^H]MODAG-005 was prepared in 30 mM Tris-HCl with 10 % ethanol (pH 7.4), to obtain a final concentration of 48 to 0.047 nM. Binding experiments were performed in triplicates for each concentration, with four independent replicates for one-hour incubation. 50 nM fibrils were incubated with decreasing concentrations for total binding and 1 µM unlabeled tracer was added to each concentration of [^3^H]MODAG-005 for non-specific binding. Plates were sealed either with a removable sealing tape (Revvity Germany Diagnostics GmbH, Lübeck, Germany) for 1 hour, to avoid evaporation and incubated at room temperature. 30 minutes prior harvesting, a glass fiber Filtermat B (Revvity Germany Diagnostics GmbH, Lübeck, Germany) was incubated in 1% polyethyleneimine (PEI). After harvesting, the filtermat was dried in a microwave, and melt-on scintillator sheets (Revvity Germany Diagnostics GmbH, Lübeck, Germany) were molten into the filtermat. After solidification at room temperature, the filter was placed in a MicroBeta sample bag (Revvity Germany Diagnostics GmbH, Lübeck, Germany) and measured using the Wallac MicroBeta® TriLux liquid scintillation counter (PerkinElmer, Waltham, MA, USA). Radioactivity was converted to bound pmol [^3^H]MODAG-005, using a calibration curve, and plotted against the fibril concentration (nmol of total fibrils). Non-linear regression analysis was performed using a one binding site analysis in GraphPad Prism (GraphPad Software, Inc., Version 10.1.1, La Jolla (CA), USA). In addition, a two binding site analysis was applied using a Scatchard plot based on the calculated specific binding. Data points for the high-affinity binding site (47 - 750 pM) and the low-affinity binding site (1.5 - 48 nM) were determined by visual inspection. Linear regression was performed for the two binding sites to determine the respective K_d_ and B_max_ values. Negative values were set to zero.

### Molecular dynamics (MD) simulations

#### Simulation protocol

The GROMACS 2022 simulation software package ^73,74^ was used to set up, carry out and analyze the MD simulations. Settings for production runs were chosen as follows: The long-range electrostatic interactions were treated using the Particle Mesh Ewald (PME) method ^75,76^. Bonds in protein and lipid molecules were constrained using the P-LINCS ^77^ algorithm. Water molecules were constrained using SETTLE ^78^ algorithm. Neighbor lists were updated with the Verlet list scheme ^74,79^. For production runs, the simulated systems were kept at a temperature of 300 K by applying the velocity-rescaling ^80^ algorithm. Initial velocities for the production runs were taken according to the Maxwell-Boltzmann distribution at 300 K. The pressure was held constant by using the Parrinello-Rahman barostat ^81^. The integration time step was set to 2 fs. The neighbor lists for non-bonded interactions were updated every 20 steps. Real-space electrostatic interactions were truncated at 1.2 nm. The van der Waals interactions were switched off between 1.0 to 1.2 nm and short-range electrostatic interactions were cut-off at 1.2 nm.

All simulations were carried out using periodic boundary conditions for the simulation box and utilized the CHARMM36m ^82,83^ protein force field together with the CHARMM-modified ^84^ TIP3P water model and the CHARMM36 lipid parameters ^85^. The OpenBabel web server ^86^ was used to create the input files for topology and parameter preparation of MODAG-005, anle138b and PiB molecules with the CHARMM General Force Field (CGenFF 4.6) ^87–90^. The topologies were converted to GROMACS format using the cgenff charmm2gmx.py script (http://mackerell.umaryland.edu/charmm_ff.shtml#gromacs).

All production simulations were preceded by a multi-step energy minimization and thermalization of the simulation systems.

#### L2 *α*Syn protofilament model with extended N- and C-terminus

To investigate small molecule binding to different parts of the lipidic polymorph L2 *α*Syn fibrils located by ssNMR measurements, we performed unbiased Molecular Dynamics (MD) simulations of an *α*Syn fibril model in the presence of POPC and POPA lipids. The cryo-EM structure of L2 fibrils (PDB ID 8A4L) was used to build a protofilament composed of 30 copies of identical peptide chains with a twist. The cryo-EM structure includes the ordered residues 14-24 and 33-96, whereas the rest of the *α*Syn sequence are not resolved. The L2 protofilament model was completed by extending the N- & C-termini by residues 10-13 (PDB ID 8ADU) and 97-100 (PDB ID 8A9L) from other structurally related *α*Syn polymorphs, such that the putative binding pocket around G93F94 is better defined. The protofilament structure was placed in a rectangular solvent box with the distance between the solute and the box edge of at least 2.5 nm. The titratable groups of the protein structure were protonated according to their standard protonation states at pH 7, and the N- and C-termini of the peptides were capped with acetyl and N-methyl groups, respectively. Counterions (Na^+^, Cl^−^) were added to yield an ionic strength of 150 mM and to neutralize the net system charge.

To relax the modeled N- and C-terminal parts, while preserving the L2 fold of the protofilament structure, we ran a 25 ns long MD simulation. Harmonic potentials to the backbone atoms of residues 14-24 and 33-96 were applied, restraining them to the initial atomic coordinates (force constant of 100 kJ mol^-1^ nm^-2^), while the rest the structure was free to move during the MD simulation. In a next step, 200 POPA and 200 POPC molecules were placed randomly in the solvent around the relaxed protofilament model structure. In addition, a MODAG-005 molecule was modeled inside the internal cavity (around Gly67Gly68) and two external binding sites (around Gly93Phe94 and Lys58-Glu61), such that their long axis aligned with the protofilament axis. The ligands were inserted without interatomic overlaps to the side-chain atoms of the surrounding residues. The final setup of the L2 *α*Syn protofilament model contained 275,000 atoms, including 62,000 water molecules. In total, 20 independent NPT production simulations (representing both possible pyrazole nitrogen protonation states), each 500 ns long, were carried out for this simulation system. Importantly, without restraining any of the atomic coordinates.

#### Model systems of putative small molecule binding sites in *α*Syn fibrils

To compare and corroborate the dynamics and energetics of small molecule binding to *α*Syn fibrils, we performed MD simulations on smaller protofilament substructures. Instead of simulating the whole fibril model, only peptide chain parts near the suspected small molecule binding sites were included. To ensure that the truncated fibril models were stable, persevering the initial fold, we restrained all backbone atoms of the residues resolved by cryo-EM to the initial atomic coordinates using a harmonic potential with a force constant of 1000 kJ mol^−1^ nm^−2^. In addition, flat-bottomed position restraints were used to keep the diffusion of the small molecules in the simulation box restricted to a cylindrical part centered on binding site of interest (radius 1 nm, force constant of 1000 kJ mol^−1^ nm^−2^). This approach reduces computational cost significantly and therefore allows to test putative multiple small molecule binding sites in L2 and other disease relevant *α*Syn fibril polymorphs with MD simulations on long time scales.

We built the following simulation systems (Supplementary Table. 6):

PDB ID 8A4L - internal binding site: residues 65-90

PDB ID 8A4L - external binding site: residues 10-24, 84-96

PDB ID 8A9L - external binding site: residues 10-24, 83-100

PDB ID 6XYQ (protofilament I) - internal binding site: residues 49-76

PDB ID 6XYQ (protofilament II) - internal binding site: residues 49-76

The fibril model systems were built in each instance from the respective cryo-EM structure coordinates and were composed of 20 copies of identical peptide chains with a twist. In case of the L2 and LBD protofilament model, the N-terminus was extended with residues 10-13 (PDB ID 8ADU) as described before. Each obtained protofilament structure was placed in a rectangular solvent box with the distance between the solute and the box edge of at least 1.5 nm. The titratable groups of the protein structure were protonated according to their standard protonation states at pH 7. In case of the MSA internal binding sites of the 6xyq fibril fold, the residue E61 was simulated in the protonated state, since it has no access to the bulk solvent. N- and C-termini of the peptides in close proximity to the binding site were capped with acetyl and N-methyl groups, respectively. Counter ions (Na^+^, Cl^−^) were added to yield an ionic strength of 150 mM and to neutralize the net system charge. MODAG-005 molecules were modeled above the internal cavities and next to the external binding sites, such that their long axis aligned with the protofilament axis. The truncated fibril models were simulated with a single MODAG-005 and four MODAG-005 molecules representing both possible pyrazole nitrogen protonation states, respectively. Supplementary table 6 summarizes the number and length of the independent NPT production simulations and configuration for each simulation system. The final setup of the partially restrained fibril models contained between 60,000 to 75,000 atoms.

Trajectory analyses were performed using tools implemented in the GROMACS program package ^74^.

#### Molecular Mechanics/Poisson Boltzmann Surface Area (MM/PBSA)

The binding energy (ΔE) of anle138b and the MODAG-005 PET tracer molecule interacting with αSyn filaments were computed using gmx MMPBSA^91^. The tool is based on AMBER’s MMPBSA.py <u>_ENREF_50</u>^92^ and able to perform end-state free energy calculations with GROMACS files.

In all instances, the data from the last 100 ns of the MD trajectories were used for MM/PBSA calculations in the single trajectory mode. Snapshots of small molecule, protein, and complex were extracted every 2500 ps from every trajectory. The binding energy was calculated from all the extracted snapshots. The average value ligand-fibril interactions and it is standard error was calculated over the results of each independent trajectory (see Supplementary Table 7).

#### Contact analysis

Short-range interatomic interactions between the MODAG-005 ligands and the fibril model were quantified with the gmx mindist program. A contact to the polar atoms of the small molecule was considered as formed, if found within a cutoff of 0.4 nm to the backbone heavy atoms of a protein residue. Contacts were averaged over time and over the individual trajectories of each simulation set. If not explicitly stated, only contacts to the core β-strand layers (neglecting the top β-strands on both protofilament edges) were considered for analysis. In addition, and according to the isotope labeling scheme in the NMR experiments, the interatomic distances between the 1,2-N atoms of the MODAG -005 molecule and the side chain aliphatic carbon atoms of protein residue Ile88, as well as the aromatic carbon atoms of protein residue Phe94 were probed. The average number of water oxygen atoms with direct contact with the MODAG-005 ligand was determined based on a 0.4 nm distance cutoff to all atoms the small molecule.

## Acknowledgments

We thank C. Schwiegk for technical help in protein expression and purification. We also thank G. Heim for the acquisition of TEM images. We thank J. Schimpfhauser, J. Bienert and V. N. Belov from the facility for Synthetic Chemistry at the Max Planck Institute for Multidisciplinary Sciences, Göttingen for synthesizing the TEMTriPol-1 radical.

## Authors contributions

S.B. oversaw protein expression and purification. AL synthesized isotope labeled MODAG-005. M.K. prepared fibril samples. AKG and DB performed and analyzed [^3^H] fibril binding assays M.K and L.B.A. performed NMR experiments. D.M. conceptualized and carried out MD-simulations. C.D. collected the cryo-EM images. B.F. processed the cryo-EM images, reconstructed the fibril structures. M.K., D.M. and B.F. prepared figures and M.K. wrote the initial draft. B.L.dG, A.G., C.G. and L.B.A. supervised the project; all authors contributed to writing the manuscript.

## Data availability

The cryo-EM maps have been deposited in the Electron Microscopy Data bank (EMDB) under the accession numbers EMD-52038 [https://www.ebi.ac.uk/pdbe/entry/emdb/ EMD-52038] (L2A-I, short), EMD-52039 [https://www.ebi.ac.uk/pdbe/entry/emdb/ EMD-52039] (L2A-II, short), EMD-52040 [https://www.ebi.ac.uk/pdbe/entry/emdb/ EMD-52040] (L2C, short), EMD-52041 [https://www.ebi.ac.uk/pdbe/entry/emdb/ EMD-52041] (L2A-I, long), EMD-52042 [https://www.ebi.ac.uk/pdbe/entry/emdb/ EMD-52042] (L2A-II, long), and EMD-52043 [https://www.ebi.ac.uk/pdbe/entry/emdb/ EMD-52043] (L3C, long). The corresponding atomic models have been deposited in the Protein Data Bank (PDB) under the accession numbers: 9hc6 [https://doi.org/10.2210/pdb9hc6/pdb] (L2A-II, short), 9hc7 [https://doi.org/10.2210/pdb9hc7/pdb] (L2A-II, short), 9hc8 [https://doi.org/10.2210/pdb9hc8/pdb] (L2C, short), 9hc9 [https://doi.org/10.2210/pdb9hc9/pdb] (L2A-I, long), 9hca [https://doi.org/10.2210/pdb9hca/pdb] (L2A-II, long), and 9hcb [https://doi.org/10.2210/pdb9hcb/pdb] (L2C, long).

Ligand topology and parameters, initial coordinate and simulation input files and a coordinate file of the final output are provided through the Edmond data repository at [https://doi.org/10.17617/3.2XY3VA].

## References

1 Stefanis, L. α-Synuclein in Parkinson’s disease. Cold Spring Harbor perspectives in medicine 2, a009399 (2012).

2 Galvin, J. E., Lee, V. M.-Y. & Trojanowski, J. Q. Synucleinopathies: clinical and pathological implications. Archives of neurology 58, 186–190 (2001).

3 Kouli, A., Torsney, K. M. & Kuan, W.-L. Parkinson’s disease: etiology, neuropathology, and pathogenesis. Exon Publications, 3–26 (2018).

4 Gibb, W., Esiri, M. & Lees, A. Clinical and pathological features of diffuse cortical Lewy body disease (Lewy body dementia). Brain 110, 1131–1153 (1987).

5 Walker, Z., Possin, K. L., Boeve, B. F. & Aarsland, D. Lewy body dementias. The Lancet 386, 1683–1697 (2015).

6 Ubhi, K., Low, P. & Masliah, E. Multiple system atrophy: a clinical and neuropathological perspective. Trends in neurosciences 34, 581–590 (2011).

7 Wenning, G. K., Colosimo, C., Geser, F. & Poewe, W. Multiple system atrophy. The Lancet Neurology 3, 93–103 (2004).

8 Cole, T. A., et al. α-Synuclein antisense oligonucleotides as a disease-modifying therapy for Parkinson’s disease. JCI insight 6 (2021).

9 Pardo-Moreno, T. et al. Current treatments and new, tentative therapies for Parkinson’s disease. Pharmaceutics 15, 770 (2023).

10 Donadio, V. et al. In vivo diagnosis of synucleinopathies: a comparative study of skin biopsy and RT-QuIC. Neurology 96, e2513–e2524 (2021).

11 Buchert, R., Buhmann, C., Apostolova, I., Meyer, P. T. & Gallinat, J. Nuclear imaging in the diagnosis of clinically uncertain parkinsonian syndromes. Deutsches Ärzteblatt International 116, 747 (2019).

12 Rong, J., Haider, A., Jeppesen, T. E., Josephson, L. & Liang, S. H. Radiochemistry for positron emission tomography. Nature communications 14, 3257 (2023).

13 Dupont, A.-C., Largeau, B., Guilloteau, D., Santiago Ribeiro, M. J. & Arlicot, N. The place of PET to assess new therapeutic effectiveness in neurodegenerative diseases. Contrast Media & Molecular Imaging 2018 (2018).

14 Tian, G. The development of [11C] M503-1619 as a PET tracer for imaging α-synucleinopathies in Parkinson’s disease (PD), Department of Radiology, University of Pittsburgh, (2023).

15 Kim, H. Y. et al. A Novel Brain PET Radiotracer for Imaging Alpha Synuclein Fibrils in Multiple System Atrophy. Journal of Medicinal Chemistry 66, 12185–12202 (2023).

16 Bennacef, I. et al. ALPHA-SYNUCLEIN BINDERS AND METHODS OF USE. WO patent WO 2024/186584 A2 (2024).

17 Borroni, E. et al. RADIOLABELED COMPOUNDS. WO patent WO 2019/121661 A1 (2019).

18 Zhu, L., Ploessl, K. & Kung, H. F. PET/SPECT imaging agents for neurodegenerative diseases. Chemical Society Reviews 43, 6683–6691 (2014).

19 Smith, R. et al. The α-synuclein PET tracer [18F] ACI-12589 distinguishes multiple system atrophy from other neurodegenerative diseases. Nature communications 14, 6750 (2023).

20 Endo, H. et al. Imaging α-synuclein pathologies in animal models and patients with Parkinson’s and related diseases. Neuron (2024).

21 Wagner, J. et al. Anle138b: a novel oligomer modulator for disease-modifying therapy of neurodegenerative diseases such as prion and Parkinson’s disease. Acta neuropathologica 125, 795–813 (2013).

22 Levin, J. et al. Safety, tolerability and pharmacokinetics of the oligomer modulator anle138b with exposure levels sufficient for therapeutic efficacy in a murine Parkinson model: A randomised, double-blind, placebo-controlled phase 1a trial. EBioMedicine 80 (2022).

23. Saw, R., et al. [11C] MODAG 005–a novel PET tracer targeting alpha-synuclein aggregates in the brain. (2024).

24 Antonschmidt, L. et al. Insights into the molecular mechanism of amyloid filament formation: Segmental folding of α-synuclein on lipid membranes. Science Advances 7, eabg2174, doi:10.1126/sciadv.abg2174 (2021).

25 Sant, V. et al. Lipidic folding pathway of α-Synuclein via a toxic oligomer. Nature Communications 16, 760 (2025).

26 Frieg, B. et al. The 3D structure of lipidic fibrils of α-synuclein. Nature communications 13, 6810 (2022).

27 Tao, Y. et al. Structural mechanism for specific binding of chemical compounds to amyloid fibrils. Nature Chemical Biology 19, 1235–1245 (2023).

28 Xiang, J. et al. Development of an α-synuclein positron emission tomography tracer for imaging synucleinopathies. Cell 186, 3350–3367. e3319 (2023).

29 Shi, Y., Ghetti, B., Goedert, M. & Scheres, S. H. Cryo-EM structures of chronic traumatic encephalopathy tau filaments with PET ligand flortaucipir. Journal of molecular biology 435, 168025 (2023).

30 Shi, Y. et al. Cryo-EM structures of tau filaments from Alzheimer’s disease with PET ligand APN-1607. Acta neuropathologica 141, 697–708 (2021).

31 Antonschmidt, L. et al. The clinical drug candidate anle138b binds in a cavity of lipidic α-synuclein fibrils. Nature communications 13, 5385 (2022).

32 Lange, A., Luca, S. & Baldus, M. Structural constraints from proton-mediated rare-spin correlation spectroscopy in rotating solids. Journal of the American Chemical Society 124, 9704–9705 (2002).

33 Takegoshi, K., Nakamura, S. & Terao, T. 13C–1H dipolar-assisted rotational resonance in magic-angle spinning NMR. Chemical physics letters 344, 631–637 (2001).

34 Sevigny, J. et al. The antibody aducanumab reduces Aβ plaques in Alzheimer’s disease. Nature 537, 50–56 (2016).

35 Klein, G. et al. Thirty-six-month amyloid positron emission tomography results show continued reduction in amyloid burden with subcutaneous gantenerumab. The journal of prevention of Alzheimer’s disease 8, 3–6 (2021).

36 Klunk, W. E. et al. Imaging brain amyloid in Alzheimer’s disease with Pittsburgh Compound-B. Annals of Neurology: Official Journal of the American Neurological Association and the Child Neurology Society 55, 306–319 (2004).

37 Jie, C. V., Treyer, V., Schibli, R. & Mu, L. Tauvid™: the first FDA-approved PET tracer for imaging tau pathology in Alzheimer’s disease. Pharmaceuticals 14, 110 (2021).

38 Barthel, H. & Sabri, O. Clinical use and utility of amyloid imaging. Journal of Nuclear Medicine 58, 1711–1717 (2017).

39 Kuebler, L. et al. [11 C] MODAG-001—towards a PET tracer targeting α-synuclein aggregates. European Journal of Nuclear Medicine and Molecular Imaging 48, 1759–1772 (2021).

40 Verdurand, M. et al. In silico, in vitro, and in vivo evaluation of new candidates for α-synuclein PET imaging. Molecular Pharmaceutics 15, 3153–3166 (2018).

41 Akasaka, T. et al. Synthesis and evaluation of novel radioiodinated phenylbenzofuranone derivatives as α-synuclein imaging probes. Bioorganic & Medicinal Chemistry Letters 64, 128679 (2022).

42 Kaide, S. et al. Chalcone analogue as new candidate for selective detection of α-synuclein pathology. ACS Chemical Neuroscience 13, 16–26 (2021).

43 Kovalska, V. et al. Tri-and pentamethine cyanine dyes for fluorescent detection of α-synuclein oligomeric aggregates. Journal of fluorescence 22, 1441–1448 (2012).

44 Neal, K. L. et al. Development and screening of contrast agents for in vivo imaging of Parkinson’s disease. Molecular Imaging and Biology 15, 585–595 (2013).

45 Korat, Š. et al. Alpha-synuclein PET tracer development—an overview about current efforts. Pharmaceuticals 14, 847 (2021).

46 Levin, J. et al. The oligomer modulator anle138b inhibits disease progression in a Parkinson mouse model even with treatment started after disease onset. Acta neuropathologica 127, 779–780 (2014).

47 Hsieh, C.-J. et al. Alpha synuclein fibrils contain multiple binding sites for small molecules. ACS Chemical Neuroscience 9, 2521–2527 (2018).

48 Kaiser, L. et al. Annual Congress of the European Association of Nuclear Medicine October 12–16, 2019 Barcelona, Spain. Eur J Nucl Med Mol Imaging 46, 1–952 (2019).

49 Matsuoka, K. et al. High-contrast imaging of α-synuclein pathologies in living patients with multiple system atrophy. Movement Disorders 37, 2159–2161 (2022).

50 Gao, M. W., M.; Glick-Wilson, B.; Meyer, J.; Peters, J.; Territo, P.; Green, M.; Hutchins, G.; Zarrinmayeh, H.; Zheng, Q.-H. Poster Presentations. Synthesis and initial in vitro characterization of [18F]fluoroalkyl derivatives of GSK1482160 as new candidate P2X7R radioligands. Journal of Labelled Compounds and Radiopharmaceuticals 62, S123–S588, 10.1002/jlcr.3725 (2019).

51 Schütz, A. K. et al. The amyloid–Congo red interface at atomic resolution. Angewandte Chemie-International Edition 50, 5956 (2011).

52 Niu, Z. et al. Mapping the Binding Interface of PET Tracer Molecules and Alzheimer Disease Aβ Fibrils by Using MAS Solid-State NMR Spectroscopy. ChemBioChem 21, 2495–2502 (2020).

53 Duan, P. et al. Binding sites of a positron emission tomography imaging agent in Alzheimer’s β-amyloid fibrils studied using 19F solid-state NMR. Journal of the American Chemical Society 144, 1416–1430 (2022).

54 Kuang, G., Murugan, N. A. & Ågren, H. Mechanistic insight into the binding profile of DCVJ and α-synuclein fibril revealed by multiscale simulations. ACS Chemical Neuroscience 10, 610–617 (2018).

55 Dervişoğlu, R. et al. Anle138b interaction in α-synuclein aggregates by dynamic nuclear polarization NMR. Methods 214, 18–27 (2023).

56 Hoyer, W. et al. Dependence of α-synuclein aggregate morphology on solution conditions. Journal of molecular biology 322, 383–393 (2002).

57 Barbet-Massin, E. et al. Rapid proton-detected NMR assignment for proteins with fast magic angle spinning. Journal of the American Chemical Society 136, 12489–12497 (2014).

58 Najbauer, E. E. & Andreas, L. B. Correcting for magnetic field drift in magic-angle spinning NMR datasets. Journal of Magnetic Resonance 305, 1–4, 10.1016/j.jmr.2019.05.005 (2019).

59 Lee, W., Tonelli, M. & Markley, J. L. NMRFAM-SPARKY: enhanced software for biomolecular NMR spectroscopy. Bioinformatics 31, 1325–1327, doi:10.1093/bioinformatics/btu830 (2014).

60 Mathies, G. et al. Efficient dynamic nuclear polarization at 800 MHz/527 GHz with trityl-nitroxide biradicals. Angewandte Chemie International Edition 54, 11770–11774 (2015).

61 Liu, Y., Villamena, F. A., Rockenbauer, A., Song, Y. & Zweier, J. L. Structural factors controlling the spin– spin exchange coupling: EPR spectroscopic studies of highly asymmetric trityl–nitroxide biradicals. Journal of the American Chemical Society 135, 2350–2356 (2013).

62 Delaglio, F. et al. NMRPipe: a multidimensional spectral processing system based on UNIX pipes. Journal of biomolecular NMR 6, 277–293 (1995).

63 Mastronarde, D. N. Automated electron microscope tomography using robust prediction of specimen movements. Journal of structural biology 152, 36–51 (2005).

64 Tegunov, D. & Cramer, P. Real-time cryo-electron microscopy data preprocessing with Warp. Nature methods 16, 1146–1152 (2019).

65 Zivanov, J., Nakane, T. & Scheres, S. H. Estimation of high-order aberrations and anisotropic magnification from cryo-EM data sets in RELION-3.1. IUCrJ 7, 253–267 (2020).

66 He, S. & Scheres, S. H. Helical reconstruction in RELION. Journal of structural biology 198, 163–176 (2017).

67 Rohou, A. & Grigorieff, N. CTFFIND4: Fast and accurate defocus estimation from electron micrographs. Journal of structural biology 192, 216–221 (2015).

68 Wagner, T. et al. SPHIRE-crYOLO is a fast and accurate fully automated particle picker for cryo-EM. Communications biology 2, 218 (2019).

69 Afonine, P. V. et al. Real-space refinement in PHENIX for cryo-EM and crystallography. Acta Crystallographica Section D: Structural Biology 74, 531–544 (2018).

70 Emsley, P. & Cowtan, K. Coot: model-building tools for molecular graphics. Acta crystallographica section D: biological crystallography 60, 2126–2132 (2004).

71 Chen, V. B. et al. MolProbity: all-atom structure validation for macromolecular crystallography. Acta crystallographica section D: biological crystallography 66, 12–21 (2010).

72 Auld, D. S. et al. Receptor binding assays for HTS and drug discovery. Assay Guidance Manual [Internet] (2018).

73 Pronk, S. et al. GROMACS 4.5: a high-throughput and highly parallel open source molecular simulation toolkit. Bioinformatics 29, 845–854 (2013).

74 Abraham, M. J. et al. GROMACS: High performance molecular simulations through multi-level parallelism from laptops to supercomputers. SoftwareX 1, 19–25 (2015).

75 Darden, T., York, D. & Pedersen, L. Particle mesh Ewald: An N⋅ log (N) method for Ewald sums in large systems. The Journal of chemical physics 98, 10089–10092 (1993).

76 Essmann, U. et al. A smooth particle mesh Ewald method. The Journal of chemical physics 103, 8577–8593 (1995).

77 Hess, B. P-LINCS: A parallel linear constraint solver for molecular simulation. Journal of chemical theory and computation 4, 116–122 (2008).

78 Miyamoto, S. & Kollman, P. A. Settle: An analytical version of the SHAKE and RATTLE algorithm for rigid water models. Journal of computational chemistry 13, 952–962 (1992).

79 Verlet, L. Computer “experiments” on classical fluids. I. Thermodynamical properties of Lennard-Jones molecules. Physical review 159, 98 (1967).

80 Bussi, G., Donadio, D. & Parrinello, M. Canonical sampling through velocity rescaling. The Journal of chemical physics 126 (2007).

81 Parrinello, M. & Rahman, A. Polymorphic transitions in single crystals: A new molecular dynamics method. Journal of Applied physics 52, 7182–7190 (1981).

82 Best, R. B. et al. Optimization of the additive CHARMM all-atom protein force field targeting improved sampling of the backbone ϕ, ψ and side-chain χ1 and χ2 dihedral angles. Journal of chemical theory and computation 8, 3257–3273 (2012).

83 Huang, J. et al. CHARMM36m: an improved force field for folded and intrinsically disordered proteins. Nature methods 14, 71–73 (2017).

84 MacKerell Jr, A. D., et al. All-atom empirical potential for molecular modeling and dynamics studies of proteins. The journal of physical chemistry B 102, 3586–3616 (1998).

85 Klauda, J. B. et al. Update of the CHARMM all-atom additive force field for lipids: validation on six lipid types. The journal of physical chemistry B 114, 7830–7843 (2010).

86 O’Boyle, N. M. et al. Open Babel: An open chemical toolbox. Journal of cheminformatics 3, 1–14 (2011).

87 Vanommeslaeghe, K. et al. CHARMM general force field: A force field for drug-like molecules compatible with the CHARMM all-atom additive biological force fields. Journal of computational chemistry 31, 671–690 (2010).

88 Vanommeslaeghe, K. & MacKerell Jr, A. D. Automation of the CHARMM General Force Field (CGenFF) I: bond perception and atom typing. Journal of chemical information and modeling 52, 3144–3154 (2012).

89 Vanommeslaeghe, K., Raman, E. P. & MacKerell Jr, A. D. Automation of the CHARMM General Force Field (CGenFF) II: assignment of bonded parameters and partial atomic charges. Journal of chemical information and modeling 52, 3155–3168 (2012).

90 Gutiérrez, I. S. et al. Parametrization of halogen bonds in the CHARMM general force field: Improved treatment of ligand–protein interactions. Bioorganic & medicinal chemistry 24, 4812–4825 (2016).

91 Valdés-Tresanco, M. S., Valdés-Tresanco, M. E., Valiente, P. A. & Moreno, E. gmx_MMPBSA: a new tool to perform end-state free energy calculations with GROMACS. Journal of chemical theory and computation 17, 6281–6291 (2021).

92 Miller III, B. R. et al. MMPBSA. py: an efficient program for end-state free energy calculations. Journal of chemical theory and computation 8, 3314–3321 (2012).

