## Supplementary Material for "Structural insight into binding of novel PET tracer MODAG-005 to lipidic *α*-Synuclein fibrils"

### Supplementary Figures

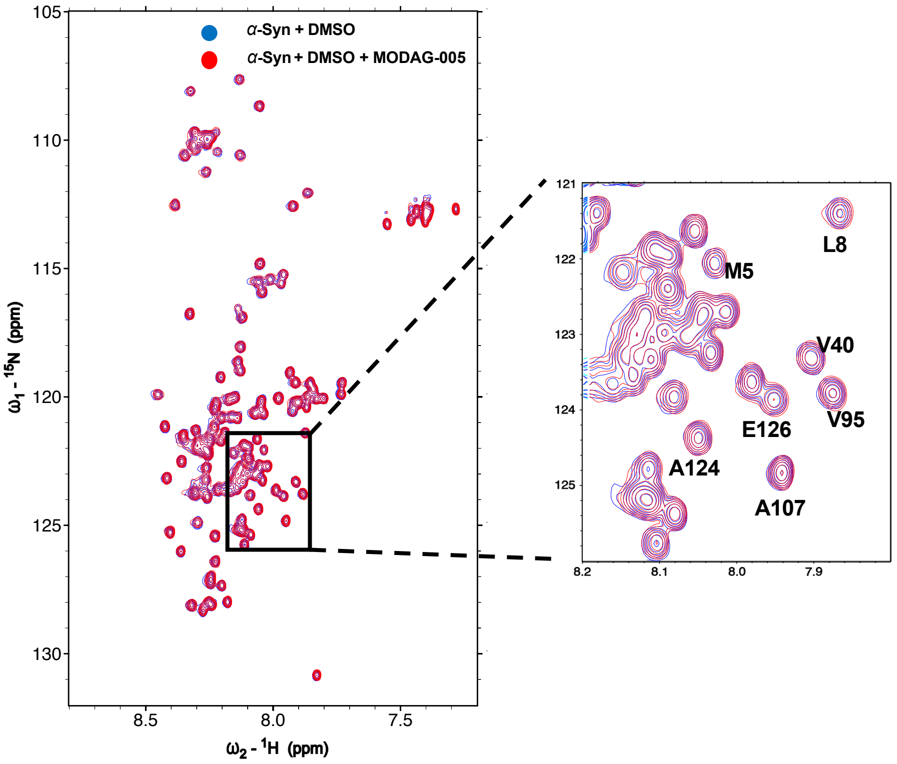


**Supplementary Fig. 1.** Effect of MODAG-005 on monomeric αSyn. Overlay of a $^{1}H$– $^{15}N$ HSQC spectra of uniformly $^{15}N$-labeled αSyn (10 μM) without MODAG-005 (blue) and with 35 μM of MODAG-005 (red). No chemical shift effects are observed and no broadening or disappearance of resonances is observed. 2% DMSO was used for these conditions.

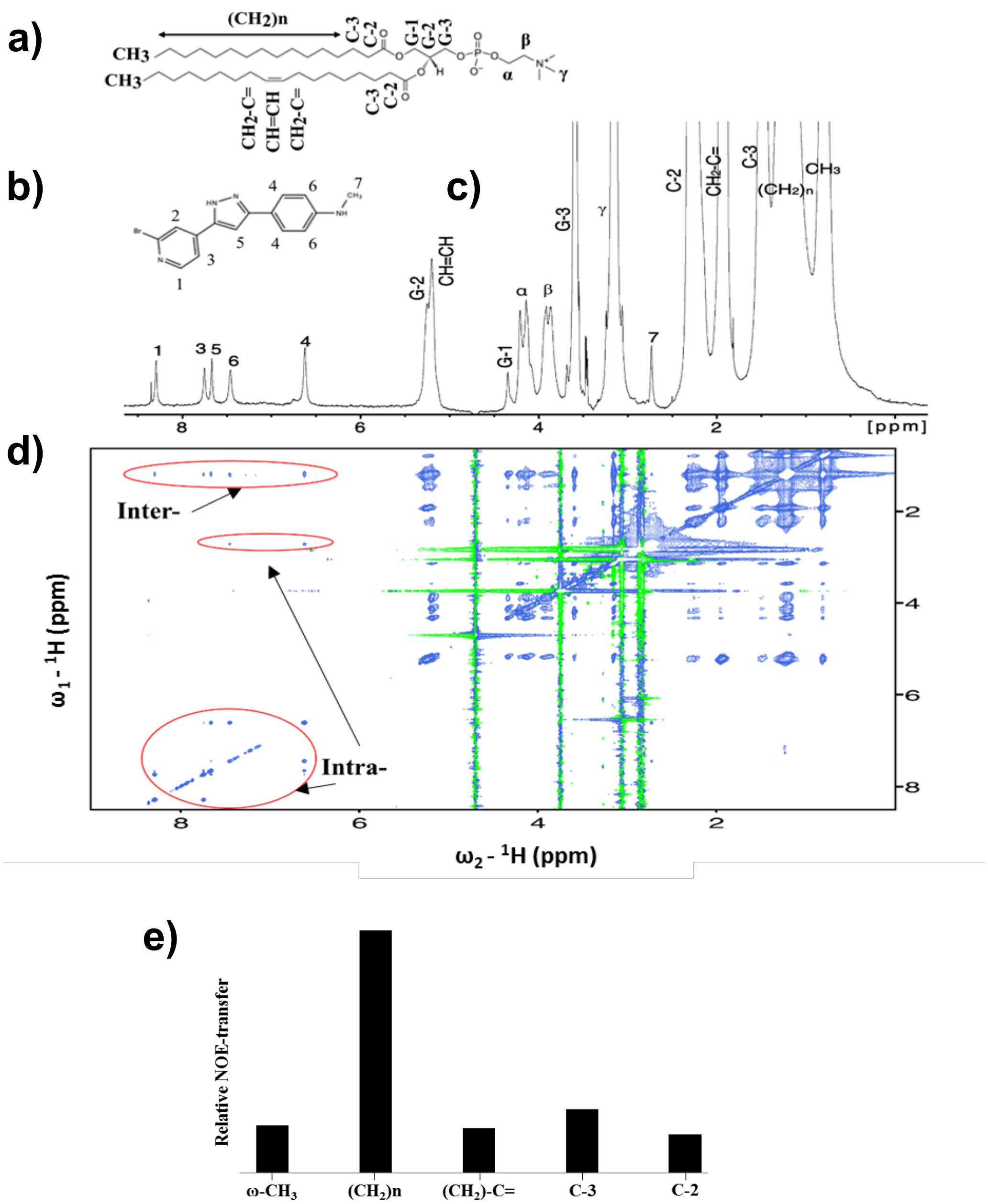


**Supplementary Fig. 2.** NOESY investigation of lipid-MODAG-005 interactions **a)** Molecular structure of POPC (1-palmitoyl-2-oleoyl-glycero-3-phosphocholine). **b)** Molecular structure of MODAG-005. **c)** 1D $^{1}H$ NMR spectra of liposome sample and assignments. **d)** $^{1}H$-$^{1}H$ NOESY Spectrum depicting inter- and intra- contacts between lipids and MODAG-005. **e)** Relative NOE-transfer rates from lipid to MODAG-005. Cross peaks between 1-2 ppm and 6-9 ppm shows intermolecular interaction between MODAG-005 and $(CH_2)n$ group providing information that the compounds are located in the middle of bilayers. A mixing time of 250 ms was used for this spectrum.

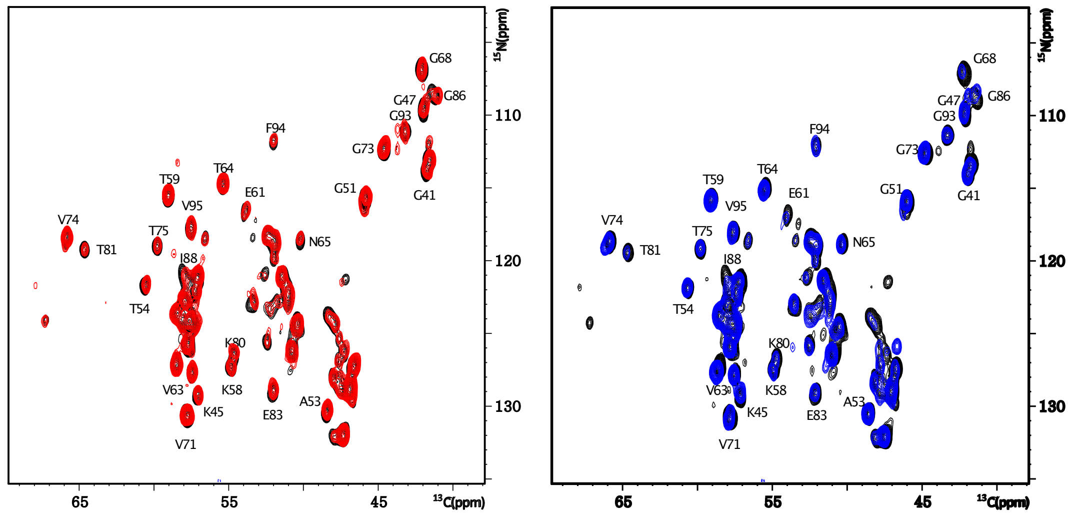


**Supplementary Fig. 3.** Overlay of the projection of hCANH spectra of L2 fibril without DMSO (black) and with the addition of 2% DMSO (red) and SUVs (blue).

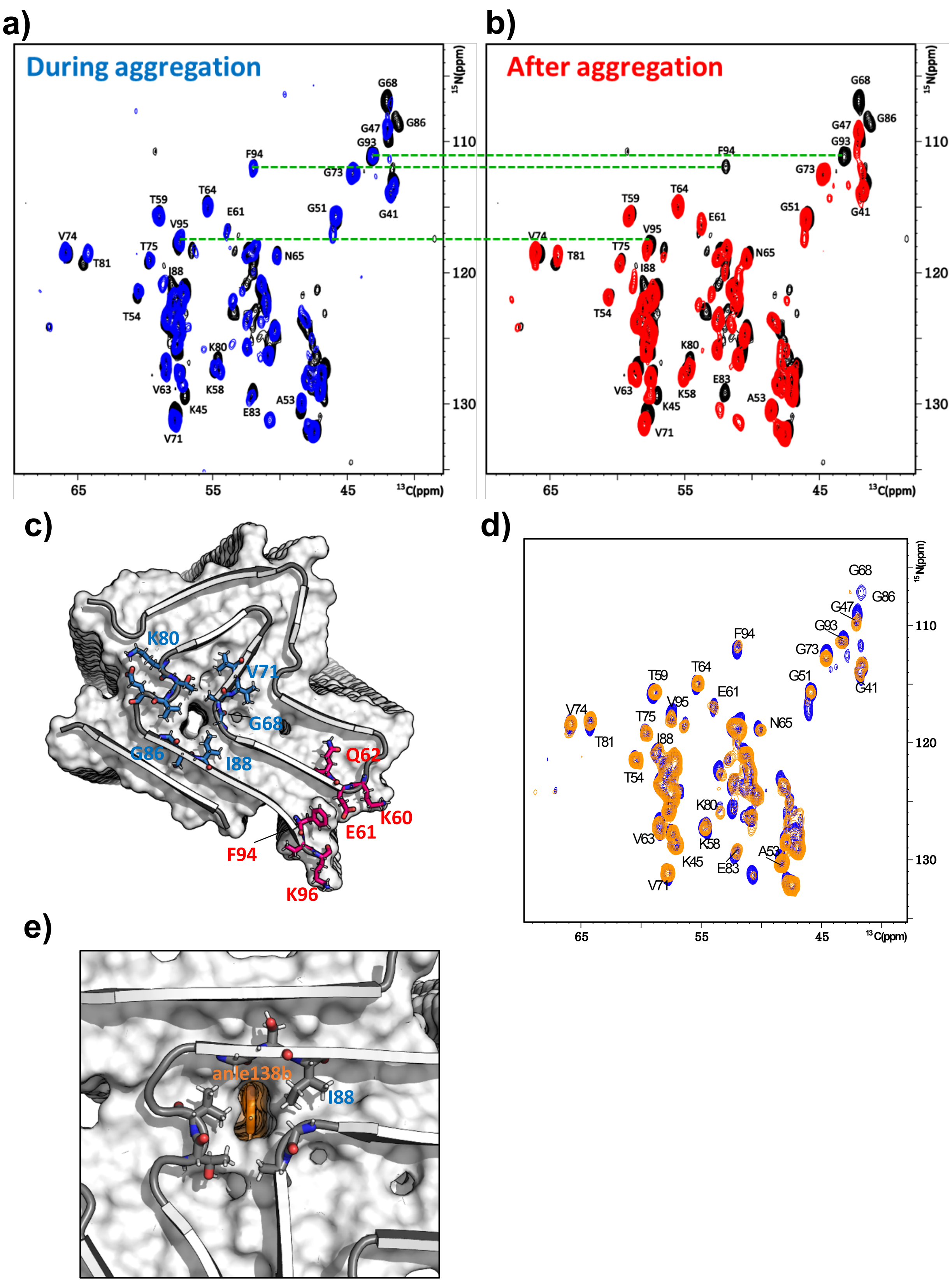
a)
During aggregation
b)
After aggregation
c)
d)
e)
$^{15}$N(ppm)
$^{13}$C(ppm)
110
120
130
65
55
45
G68
G86
G47
G93
F94
G73
G41
G51
T59
T64
E61
V95
V74
T81
T75
N65
I88
T54
K80
V63
K58
K45
E83
A53
V71
K80
V71
G68
G86
I88
Q62
K60
E61
F94
K96
anle138b
I88

**Supplementary Fig. 4.** Overlay of the projection of hCANH spectra. L2 without MODAG-005 (black) and the addition of MODAG-005 in vesicles **a)** during aggregation (blue) and **b)** in DMSO after aggregation (red). **c)** Residues that are affected by MODAG-005 with the same color coding as used in **a)** and **b)**. **d)** Overlay spectra of L2 with MODAG-005 (blue) and anle138b (orange) in the presence of vesicles in during aggregation. **e)** anle138b binding site in the L2 inner cavity. Dashed green lines indicate residues that are influenced differently depending on the timing of administration.

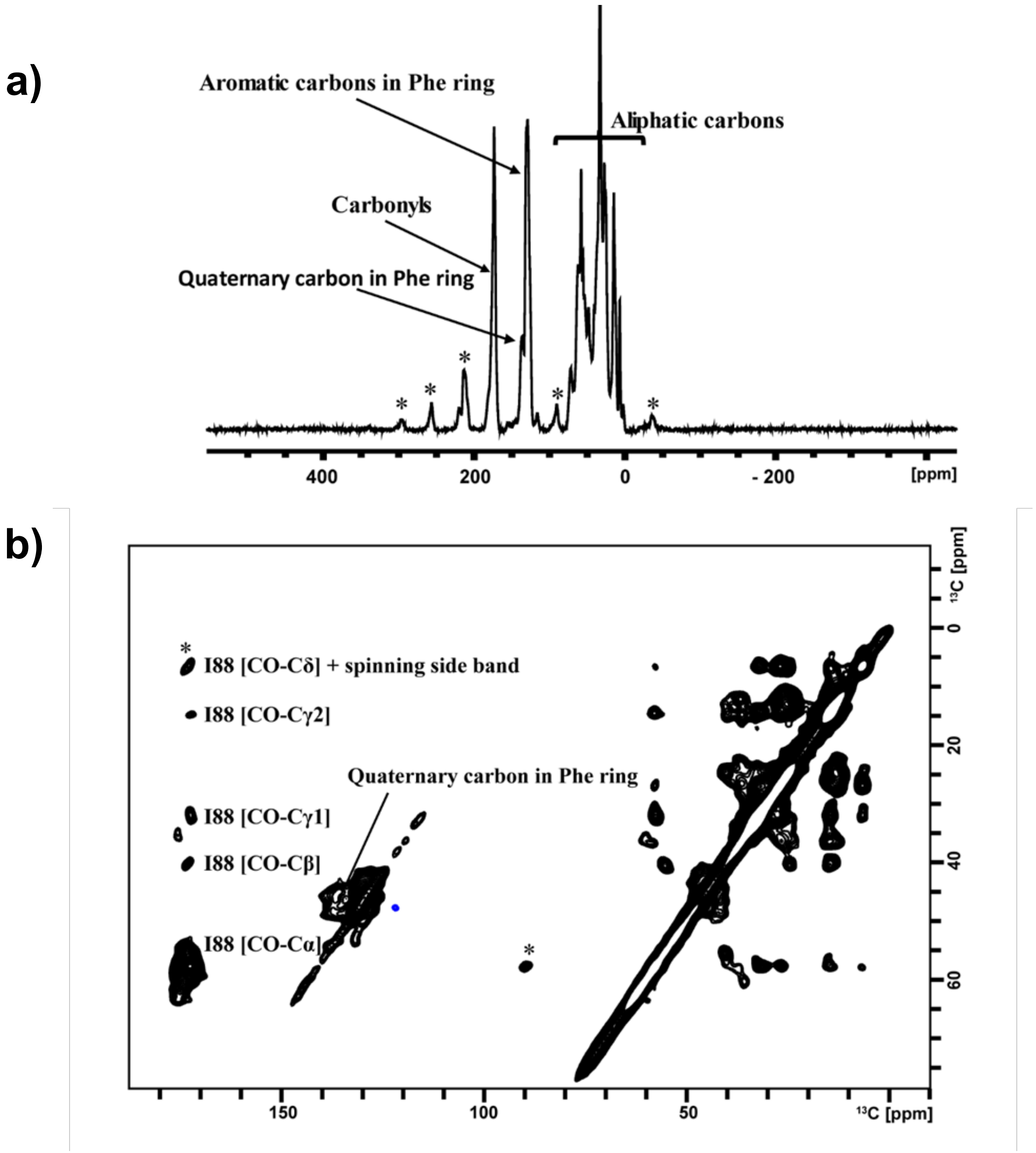


**Supplementary Fig. 5.** DNP-enhanced ssNMR of L2 fibrils of αSyn labeled with $^{13}$C-$^{15}$N Ile and $^{13}$C ring labeled Phe prepared in the presence of $^{15}$N-pyrazole nitrogen-labeled MODAG-005 dissolved in phospholipids. **a)** 1D $^{1}$H-$^{13}$C- CP-MAS spectra under DNP conditions with microwave ON **b)** $^{13}$C-$^{13}$C 2D correlation spectrum of the aggregates (L2) with $^{15}$N labeled MODAG-005. The RFDR mixing time was 1.3 ms, the MAS 12.5 kHz, and the sample temperature 100 K (DNP conditions). (*) denotes spinning sidebands.

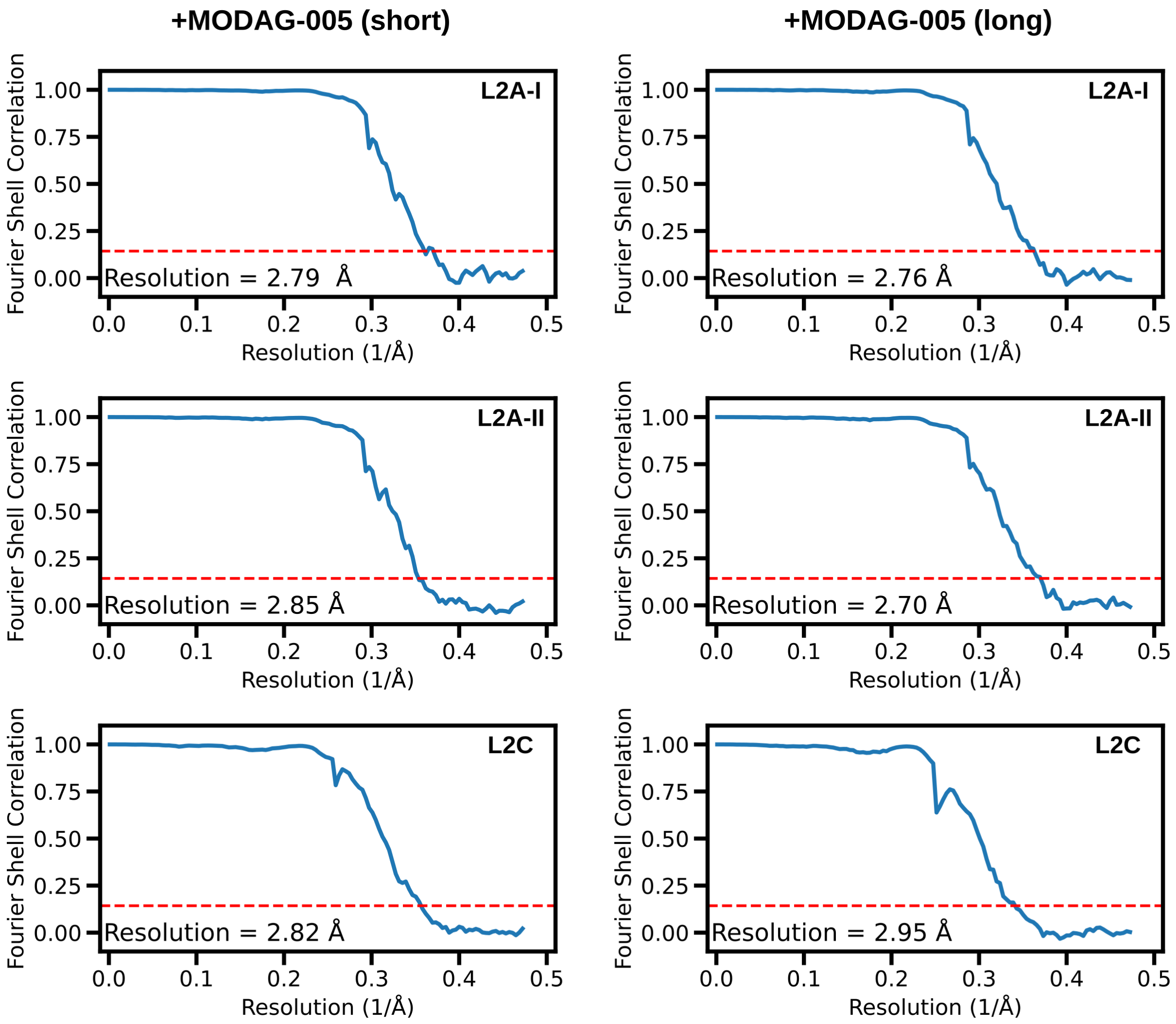


**Supplementary Fig. 6.** Fourier shell correlation curves. Masked-corrected (z-percentage is 0.1) Fourier shell correlation (FSC) curves. The final resolution is shown in the plot and was estimated from the value of the FSC curve for two separately refined masked half-maps at 0.143 (red line).

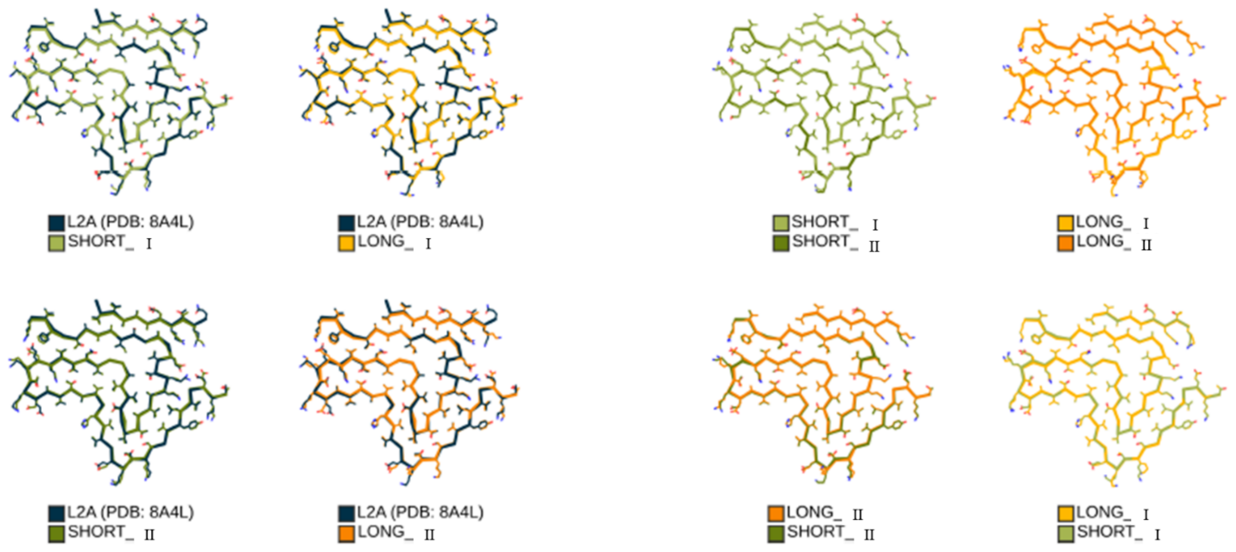


**Supplementary Fig. 7.** Comparison of cryo-EM structure of aggregates (L2A) containing MODAG-005 depending on the incubation time. **a)** Overlay of L2A structure with Short_Ⅰ/Ⅱ and Long_Ⅰ/Ⅱ indicate that addition of MODAG-005 in DMSO doesn't change the frame of L2A. **b)** Comparison between Short_Ⅰ/Ⅱand Long_Ⅰ/Ⅱ shows two distinct populations of _Ⅰ and Ⅱ have same structure. Color code: L2A (blue), Short_Ⅰ (light green), Short_Ⅱ (dark green), Long_Ⅰ (yellow) and Long_Ⅱ (orange).

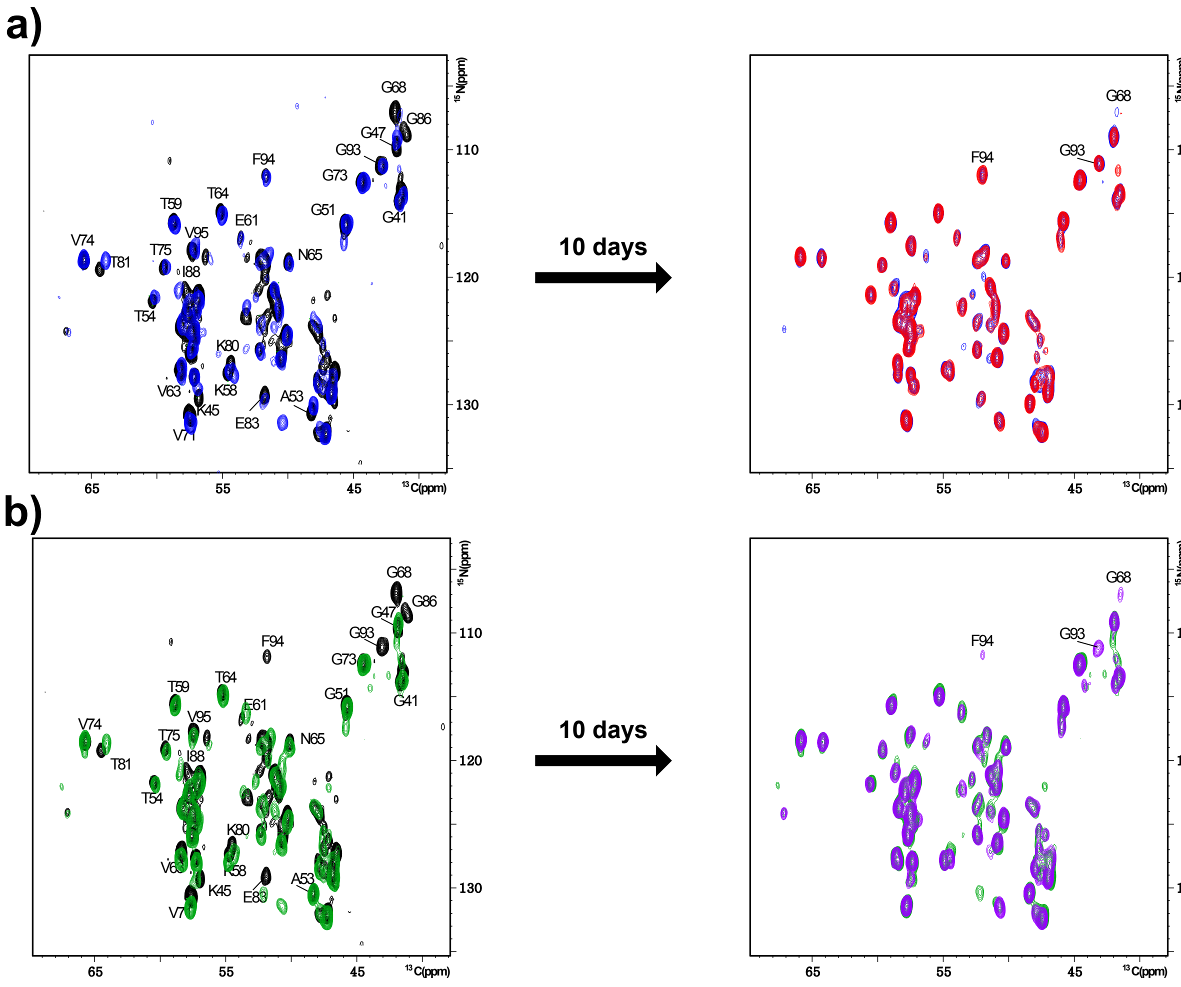


**Supplementary Fig. 8.** Comparison of the hCANH spectra of L2 prepared with MODAG-005 in vesicles during aggregation and with MODAG-005 added in DMSO after aggregation was finished. Spectra obtained directly and after ten days are compared. **a)** Overlay of the spectra of L2 in the absence of MODAG-005 (black) and in the presence (MODAG-005 in vesicles during aggregation) (blue), highlighting the changes observed after ten days period (red). **b)** Overlay of the spectra of L2 in the absence of MODAG-005 (black) and with (MODAG-005 added in DMSO after aggregation was finished) (green), highlighting the changes observed after ten days period (purple).

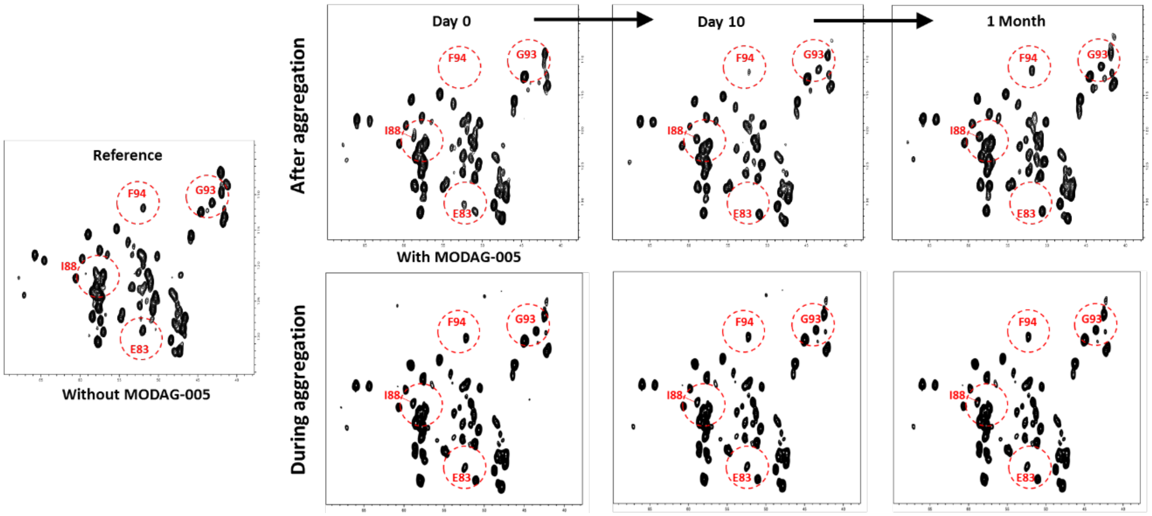


**Supplementary Fig. 9.** Comparison of hCANH spectra of αSyn aggregates (L2) over different time intervals with MODAG-005 added in DMSO to fibrils (top). Bottom: same as top but MODAG-005 was added in liposomes during aggregation. Adding MODAG-005 in DMSO after aggregation with long incubation is similar to adding MODAG-005 in liposomes during aggregation. Day 0 = 1h.

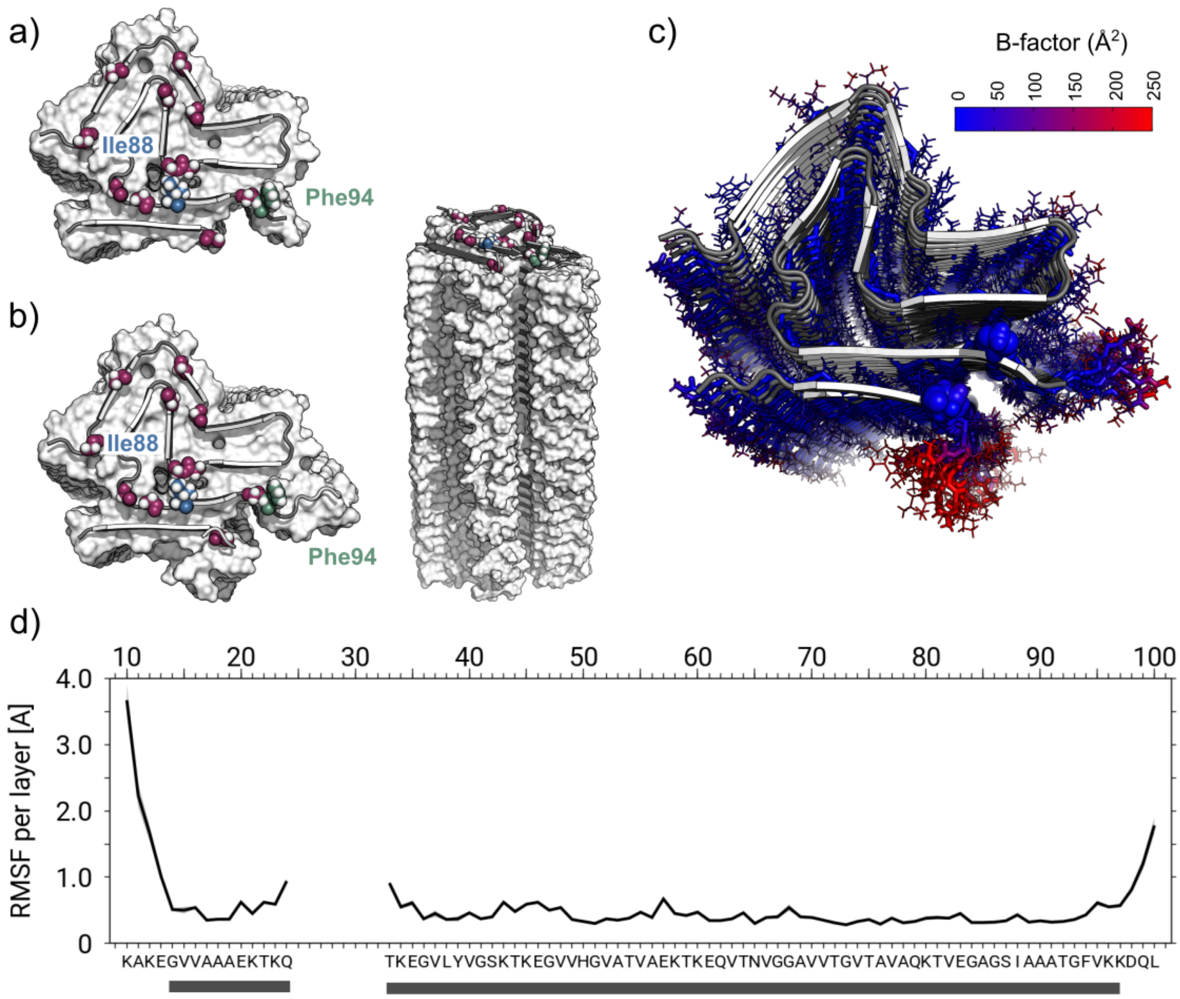


**Supplementary Fig. 10.** Simulation of the αSyn protofilament L2 with N- and C-terminus extended beyond the rigid part according to cryo-EM. **a)** The cryo-EM structure of the αSyn protofilament L2 (residues 14-24, 33-96) and **b)** simulation model with the N- and C-terminal extension of L2 (residues 10-24 i.e. addition of residues 10 to 13, 33-100 i.e. addition of residues 97-100) is shown in surface representation and with colored coded residues Gly (magenta), Ile88 (blue), and Phe94 (green) are shown as spheres. The full model of L2 with N- and C-terminal extension is also shown from the side in surface representation with all 30 stacked and twisted β-strand layers. **c)** Cartoon and stick representation of L2 with sequence extension viewed from the top. Residues G14 and G93 are shown by spheres. Sticks are color-coded according to the B-factor for each atom observed during the simulations. **d)** The root mean squared fluctuation (RMSF) for each residue (backbone heavy atoms only) are reported over all β-strand layers. Residues labeled with gray bar denoted the residues which are restrained to their starting coordinates with harmonic potentials. Data are presented as mean values ± SD (depicted by shading).

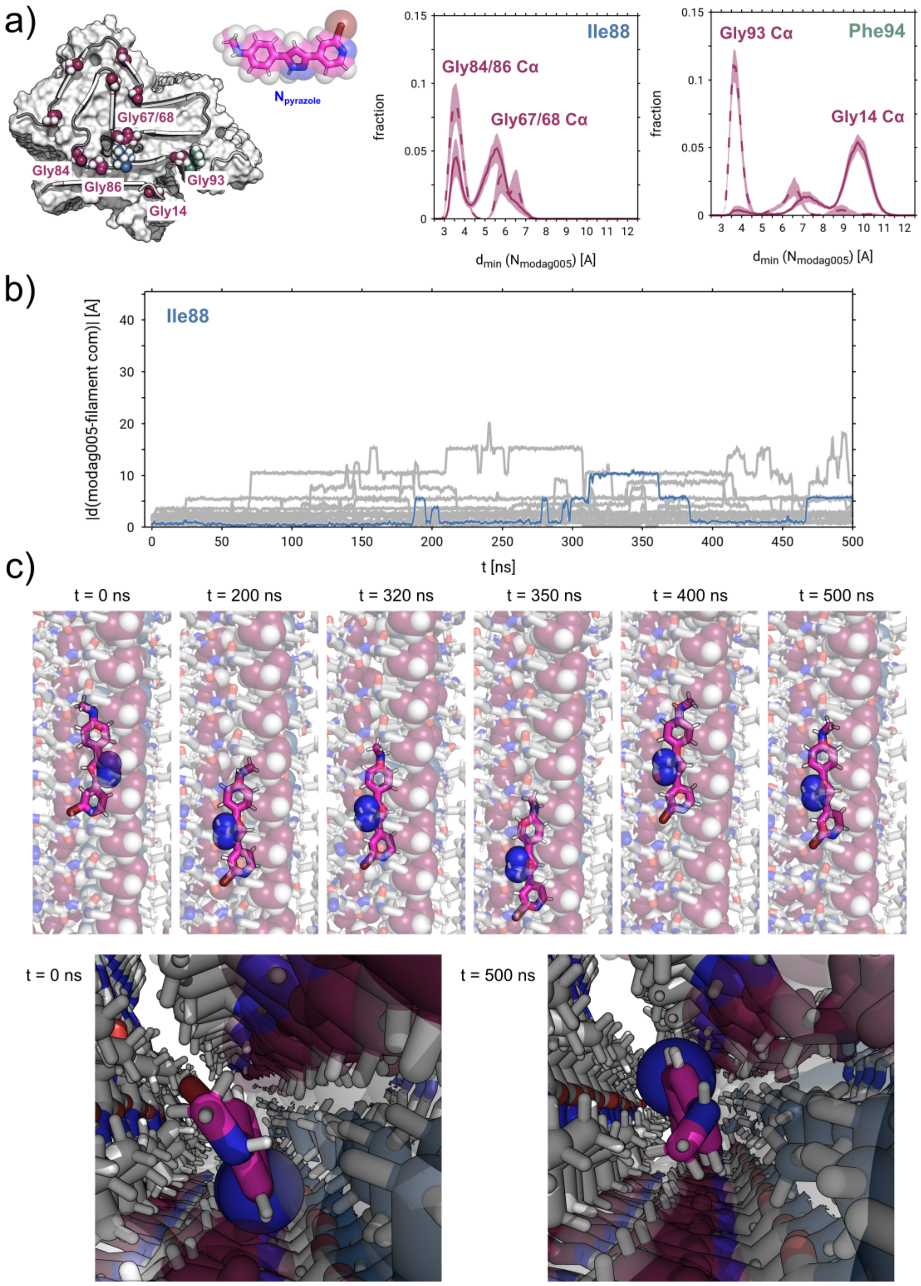


**Supplementary Fig. 11.** MODAG-005 binds dynamically along the protofilament axis in the internal and external tubular cavity of L2 αSyn fibrils. **a)** The cryo-EM structure derived model of L2 with the N- and C-terminal extension (residues 10-24, 33-100) is shown in surface representation with color-coded residues Gly (magenta), Ile88 (blue), and Phe94 (green) as spheres. Distributions of minimal distances between pyrazole nitrogen atoms of MODAG-005 and the $C_{\alpha}$ of G67, G68 and G84, G86 (broken lines) for the internal binding site. Distributions of minimal distances between pyrazole nitrogen atoms of MODAG-005 and the $C_{\alpha}$ of G14 and G93 (broken lines) for the external binding site. **b)** Distance of the Ile88 in the fibril center-of-mass to the pyrazole nitrogen atoms of MODAG-005 (representative trajectory as bold lines; others as shaded thin lines for clarity) and **c)** snapshots from MD simulations of MODAG-005 in the internal binding site: Discrete translational motion along the protofilament axis is shown. Typical alignment and multiple binding poses of MODAG-005 molecules with individual β-strand layers in the internal binding site during the MD simulations.

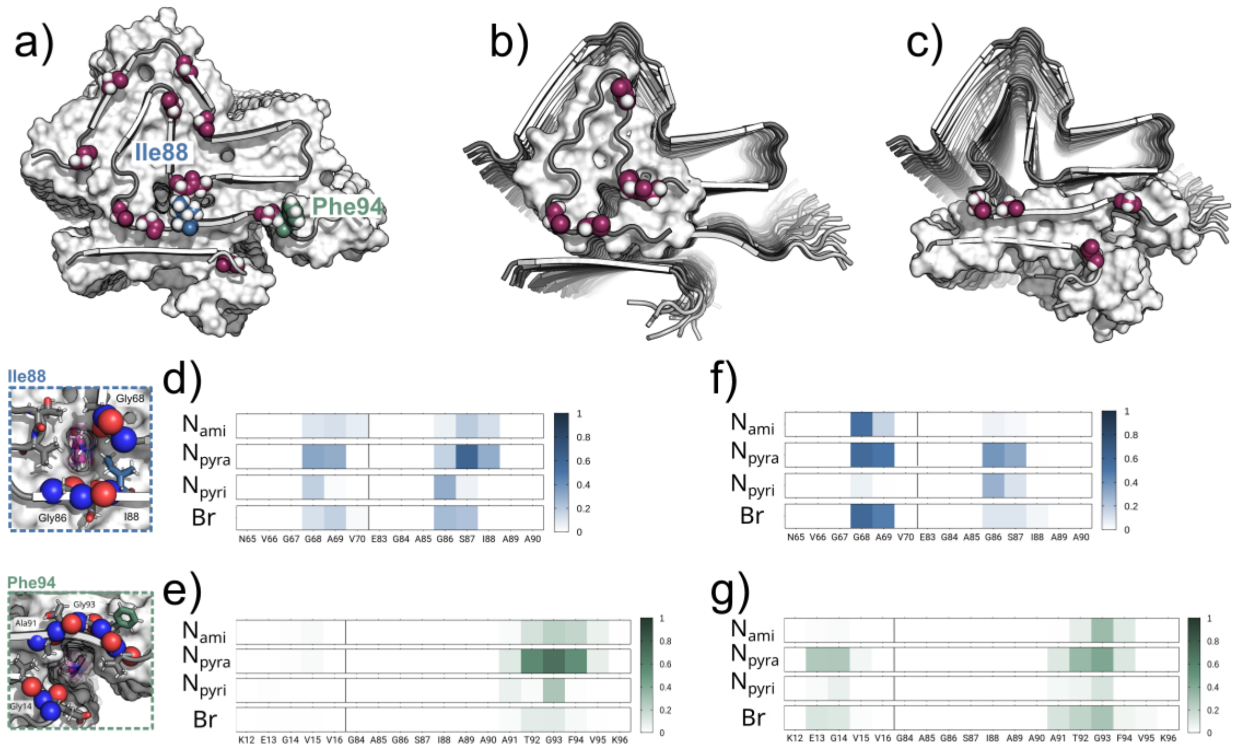


**Supplementary Fig. 12.** Comparison of polar interaction patterns with the protein backbone for full-length and truncated L2 simulation systems **a)** Starting structure for MD simulations of the N- and C-terminally extended L2 simulation fibril model. The truncated model systems centered around the respective location of the two putative MODAG-005 binding sites: **b)** The internal cavity (light blue box) of αSyn protofilament L2 between residues $_{67}$GGAV$_{70}$ and $_{80}$KTVEGAGSI$_{88}$ and **c)** the external cavity (light green box) between residues $_{14}$GVV$_{16}$ and $_{91}$ATGFV$_{95}$ (shown without MODAG-005 for clarity). Gly (magenta), Ile88 (blue), and Phe94 (green) residues are shown as color coded spheres. The truncated αSyn protofilament L2 simulation models (residues 65-90 and residues 10-24, 83-100) are shown in surface representation with the N- and C-terminally extended simulation model of L2 shown as cartoon in the background. Below, tubular cavities of internal and external binding site with MODAG-005 bound are shown, viewed up close and down the long axis of the αSyn protofilament. Polar contacts between individual residues (backbone atoms only) and polar moieties of MODAG-005 in simulations of internal (light blue; **d)** – full-length, extended model, **f)** – truncated model) and external (light green; **e)** – full-length, extended model, **g)** – truncated model) binding poses. Scale bars (right) indicate contact probabilities.

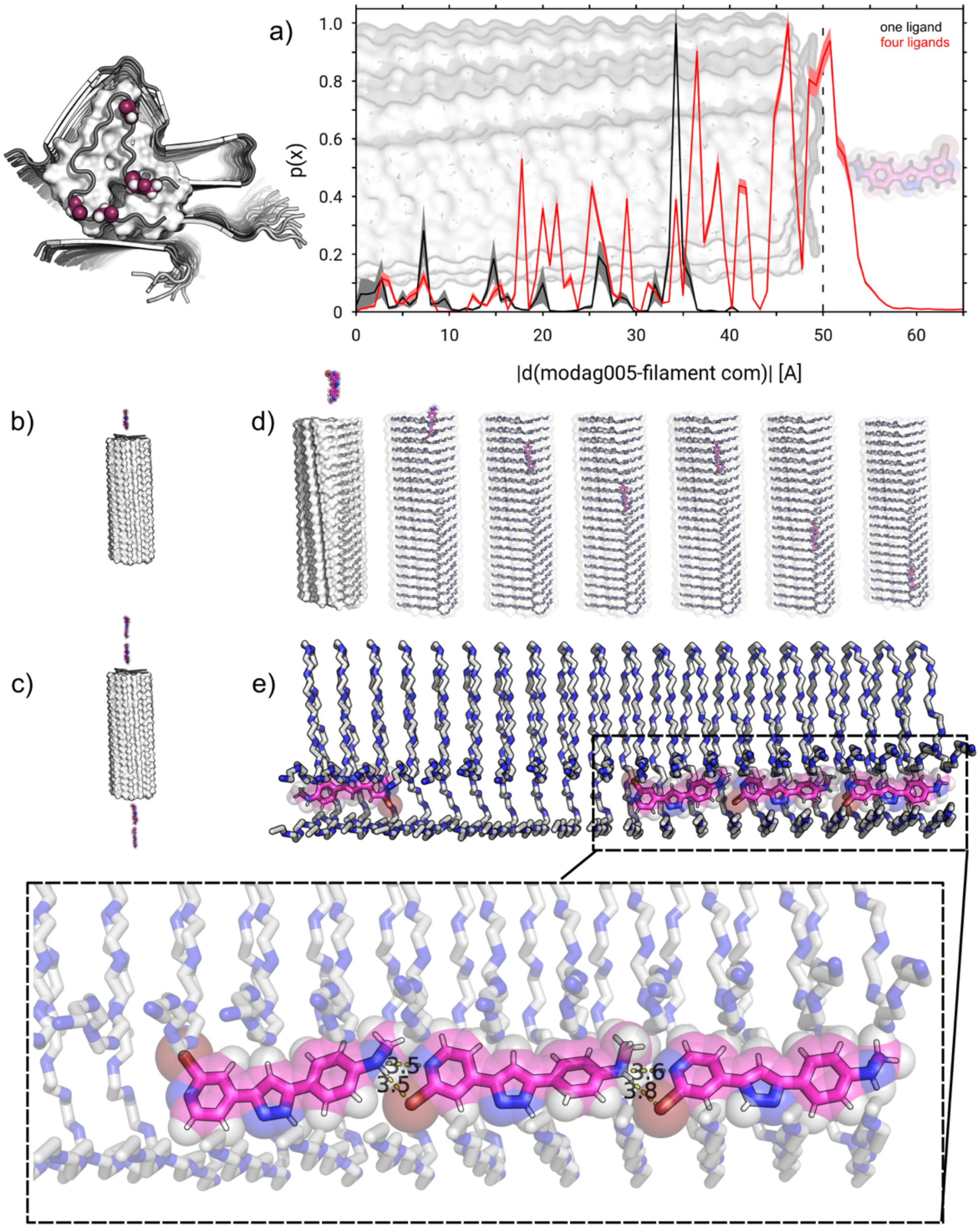


**Supplementary Fig. 13.** MODAG-005 binds spontaneously and with high affinity to the internal tubular cavity of the L2 αSyn fibril protofilament. **a)** Binding profiles *p(x)* as a function of MODAG-005 insertion depth *x* for combined simulation sets of the truncated model system centered around the internal cavity of αSyn protofilament L2 (see Fig. S12) starting with **b)** one (black lines) and **c** four copies (red lines) of MODAG-005 outside the protofilament, respectively. Data are presented as mean values ± SEM (depicted by shading). **d)** A side view of representative snapshots illustrates the spontaneous binding of MODAG-005 to the internal tubular binding site and the ligand's translational motion inside the αSyn L2 protofilament structure on the µs simulation time scale. **e)** Final snapshot (t = 1 µs) of a representative MD simulation showing all four copies of the MODAG-005 compound vertically bound inside the internal cavity of the protofilament. Enlarged view of vertical stack of three MODAG-005 molecules highlights close polar inter-ligand distances (given in Å).

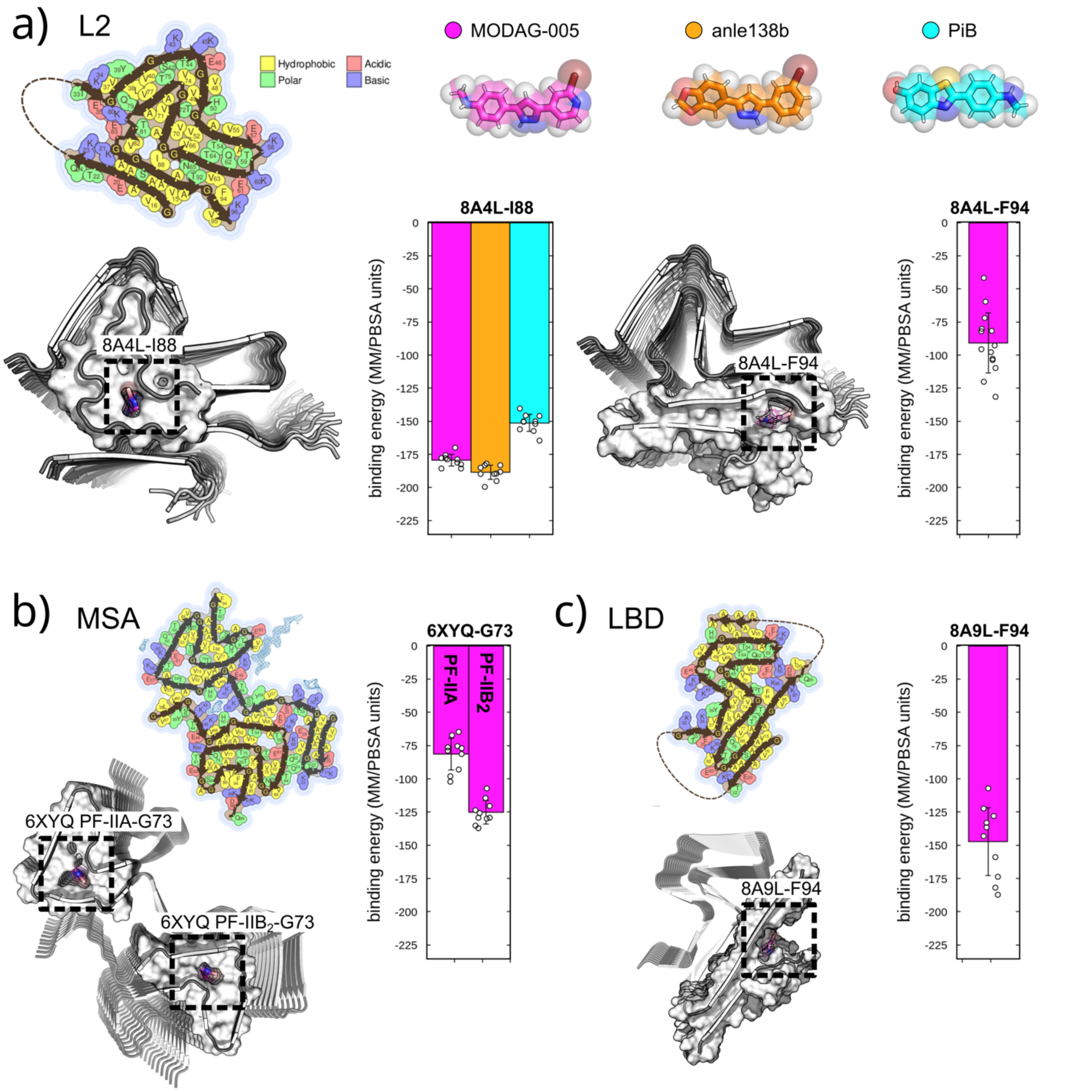


**Supplementary Fig. 14.** Energetic analysis of MODAG-005 and anle138b binding to tubular cavities of L2, MSA and LBD αSyn fibril structures. Space-filling illustrations as reported in the Amyloid Atlas 2023[1] of αSyn cryo-EM protofilaments: **a)** recombinant αSyn polymorph L2 (PDB ID: 8A4L), **b)** MSA ex vivo fibril (protofilament IIA, protofilament $IIB_2$; PDB ID: 6XYQ) and **c)** fold of PD/LBD ex vivo fibril (PDB ID: 8A9L). Tubular fibril cavities as putative small molecule - MODAG-005, anle138b and Pittsburgh Compound B (PiB) - binding sites in the respective structures are highlighted below. Binding energies as obtained from MMBPSA calculations from snapshots of independent MD simulation trajectories for MODAG-005 binding to truncated fibril models (magenta; L2 – internal, L2 - external, MSA and PD/LBD), anle138b (orange; L2 – internal) and PiB (cyan; L2 – internal) are reported as mean values ± SD. We note that due to its relative simplicity, MM/PBSA calculations offer an effective and efficient means to compute and compare binding energies of protein–ligand complexes, however, they do not allow for a quantitative prediction of experimental binding affinities.

**Supplementary Table. 1.** Average liposomes size from dynamic light scattering (DLS). Abbreviations: D (Diffusion coefficient), R(Radius), Diam (Diameter), Pd (Polydispersity), MW-R (Estimated molar mass) and Int (Intensity).

| Item | D(cm²/s) | R(nm) | Diam(nm) | Pd(nm) | %Pd | MW-R(kDa) | %Int | %Mass |
|---|---|---|---|---|---|---|---|---|
| Without MODAG-005 | 9.80647e-008 | 24.5688 | 49.1376 | 14.7874 | 60.1877 | 6031.15 | 100 | 100 |
| With MODAG-005 | 8.98272e-008 | 26.8219 | 53.6438 | 16.9515 | 63.2001 | 7405.57 | 100 | 100 |

**Supplementary Table. 2.** List of the samples for hCANH experiment.

| Sample # | Administration of MODAG-005 | Protein lab. | MODAG-005 lab. | Lipid/ Protein ratio | Incubation time | DMSO (%) | Rotor type (Sapphire/Zirconia) | MAS (kHz) | Magnet |
|---|---|---|---|---|---|---|---|---|---|
| 1 | During aggregation with lipid | $^{2}H$-$^{13}C$-$^{15}N$ | - | 10 | 1 hour, 10 days, 30 days | 0 | zirconia | 55 | 800 |
| 2 | After aggregation with lipid | $^{2}H$-$^{13}C$-$^{15}N$ | - | 10 | 1 hour, 10 days | 0 | zirconia | 55 | 800 |
| 3 | After aggregation without lipid | $^{2}H$-$^{13}C$-$^{15}N$ | - | 10 | 1 hour, 10 days, 30 days | 2 | zirconia | 55 | 800 |

**Supplementary Table. 3.** List of the samples for DNP experiment.

| Sample # | Protein lab. | MODAG-005 lab. | Lipid/ Protein ratio | Radical | Concentration of radical | Glycerol % | Rotor type (Sapphire/Zirconia) | MAS (kHz) | Magnet |
|---|---|---|---|---|---|---|---|---|---|
| 1 | U-$^{13}C$ | $^{15}N$ | 10 | TEMTriPol-1 | 8mM | 50% | zirconia | 12.5 | 600 |
| 2 | $^{13}C$-$^{15}N$ Ile, $^{13}C$ Phe ring | $^{15}N$ | 10 | TEMTriPol-1 | 8mM | 50% | zirconia | 12.5 | 600 |
| 3 | $^{13}C$-$^{15}N$ Phe | $^{15}N$ | 10 | TEMTriPol-1 | 8mM | 50% | zirconia | 12.5 | 600 |

### Supplementary Table 4 | Cryo-EM structure determination statistics.

| | Short incubation with MODAG-005 | | | Long incubation with MODAG-005 | | |
|---|---|---|---|---|---|---|
| Microscope | Titan Krios G2 | | | Titan Krios G2 | | |
| Voltage [keV] | 300 | | | 300 | | |
| Detector | K3 | | | K3 | | |
| Magnification | 81,000 | | | 81,000 | | |
| Pixel size [Å] | 1.05 | | | 1.05 | | |
| Defocus range [µm] | -0.6 to -2.3 | | | -0.6 to -2.3 | | |
| Exposure time [s/frame] | 2.85 | | | 2.85 | | |
| Number of frames | 40 | | | 40 | | |
| Total dose [$e^-/Å^2$] | 40 (1.0 $e^-/Å^2$/frame) | | | 40 (1.0 $e^-/Å^2$/frame) | | |
| **Reconstruction** | | | | | | |
| Micrographs | 17,071 | | | 15,828 | | |
| Box width [pixels] | 250 | | | 250 | | |
| Inter-box distance [pixels] | 14 | | | 14 | | |
| Picked segments (no.) | 8,928,869 | | | 10,009,818 | | |
| | **L2A-I** | **L2A-II** | **L2C** | **L2A-I** | **L2A-II** | **L2C** |
| PDB-ID | 9hc6 | 9hc7 | 9hc8 | 9hc9 | 9hca | 9hcb |
| EMDB-ID | EMD-52038 | EMD-52039 | EMD-52040 | EMD-52041 | EMD-52042 | EMD-52043 |
| Final segments [no.] | 200,200 | 130,611 | 185,538 | 129,462 | 108,610 | 149,095 |
| Final resolution [Å] (FSC=0.143) | 2.79 | 2.85 | 2.82 | 2.76 | 2.70 | 2.95 |
| Applied map sharpening B-factor [$Å^2$] | -119 | -121 | -109 | -119 | -109 | -99 |
| Symmetry imposed | C3 | C3 | C1 | C3 | C3 | C1 |
| Helical rise [Å] | 4.69 | 4.70 | 4.68 | 4.70 | 4.70 | 4.68 |
| Helical twist [°] | -0.74 | -0.60 | -0.78 | -0.73 | -0.60 | -0.77 |

**Supplementary Table 5 | Model building statistics.**

| | Short incubation with MODAG-005 | | | Long incubation with MODAG-005 | | |
|---|---|---|---|---|---|---|
| | **L2A-I** | **L2A-II** | **L2C** | **L2A-I** | **L2A-II** | **L2C** |
| **Initial model [PDB code]** | 8a4l | 8a4l | 8a4l | 8a4l | 8a4l | 8a4l |
| **Model composition** | | | | | | |
| Chains | 15 | 15 | 10 | 15 | 15 | 10 |
| Non-hydrogen atoms | 7,560 | 7,560 | 5,040 | 7,560 | 7,560 | 5,040 |
| Protein residues | 1,095 | 1,095 | 730 | 1,095 | 1,095 | 730 |
| **RMS deviations** | | | | | | |
| Bond lengths [Å] | 0.01 | 0.02 | 0.02 | 0.01 | 0.02 | 0.02 |
| Bond angles [°] | 2.03 | 2.14 | 2.00 | 2.03 | 2.12 | 2.2 |
| **Validation** | | | | | | |
| MolProbity score | 1.83 | 1.78 | 1.55 | 1.86 | 1.76 | 2.3 |
| Clashscore | 14.64 | 12.85 | 10.69 | 15.61 | 12.27 | 13.78 |
| **Ramachandran plot** | | | | | | |
| Outliers [%] | 0 | 0 | 0 | 0 | 0 | 0 |
| Allowed [%] | 2.9 | 2.9 | 1.5 | 2.9 | 2.9 | 2.9 |
| Favored [%] | 97.1 | 97.1 | 98.5 | 97.1 | 97.1 | 97.1 |

**Supplementary Table. 6.** Summary of simulation data, listing the number of independent MD simulation replica and cumulative length of the trajectories [time in µs].

| System | Binding site (PDB ID) | Ligand configuration per binding site | No. of independent simulations | Total simulation time ($\mu$s) |
|---|---|---|---|---|
| $\alpha$Syn$_{10-24,33-100}$ | internal & external in full length L2 (8A4L) | one MODAG-005 | 20 | 20 |
| $\alpha$Syn$_{65-90}$ | internal L2 (8A4L) | one MODAG-005 | 10 | 10 |
| | internal L2 (8A4L) | four MODAG-005 | 5 | 5 |
| | internal L2 (8A4L) | one anle138b | 10 | 10 |
| | internal L2 (8A4L) | one PiB | 10 | 10 |
| $\alpha$Syn$_{10-24,84-96}$ | external L2 (8A4L) | one MODAG-005 | 10 | 10 |
| | external L2 (8A4L) | four modag005 | 5 | 5 |
| $\alpha$Syn$_{10-24,83-100}$ | external LBD (8A9L) | one MODAG-005 | 10 | 10 |
| $\alpha$Syn$_{49-76}$ | internal MSA (6XYQ I) | one modag005 | 10 | 10 |
| | internal MSA (6XYQ II) | one modag005 | 10 | 10 |

**Supplementary Table. 7.** Energies of MODAG-005, anle138b and PiB binding to tubular cavities of L2, MSA and LBD αSyn fibril structures obtained from Molecular Mechanics/Poisson Boltzmann Surface Area (MM/PBSA) calculations) are reported as mean values ± SD.

| **System** | **Binding energy (MM/PBSA units)** |
|---|---|
| MODAG-005 - internal L2 (8A4L) | -180.44 +- 9.99 |
| anle138b - internal L2 (8A4L) | -188.54 +/- 5.35 |
| PiB - internal L2 (8A4L) | -151.23 +/- 6.48 |
| MODAG-005 - external L2 (8A4L) | -94.41 +/- 20.22 |
| MODAG-005 - internal MSA PF-IIA (6XYQ) | -81.44 +/- 11.87 |
| MODAG-005 - internal MSA PF-IIB2 (6XYQ) | -125.21 +/- 8.70 |
| MODAG-005 - external LBD (8A9L) | -151.23 +/- 6.48 |
